# Moth species richness in an upland tropical rainforest: A citizen scientist assisted study

**DOI:** 10.1101/2021.11.07.467659

**Authors:** David Y. P. Tng, Deborah M. G. Apgaua, Nicholas J. Fisher, Victor W. Fazio

## Abstract

Diversity studies on moths in Australia are rare, presenting various shortfalls in knowledge that impedes and understanding of their biodiversity values and their conservation. In particular, the Wet Tropics of Australia deserves attention, given the paucity of systematic moth surveys in the region and its World Heritage Area status. To fill this knowledge gap, we conducted a study to observe moths on 191 nights over a main one-year survey period at an upland rainforest locality, and uploaded all observations on iNaturalist. We also compiled other incidental observations in the general locality by other observers and observations outside the survey period. In total, we document 4,434 observations of moths represented by 1041 distinct moth morphospecies. Of these, 703 are formally named species of moths, 146 to genus and 255 to higher taxonomic designations above genus level. Despite the rather intensive main survey effort, our results suggest that we have yet to reach a plateau in documenting the moth species richness of the locality. Using this study as a model, we show that the iNaturalist platform serves as an effective means to document and digitally curate biodiversity values at a locality, whilst providing complete data transparency and enabling broader community engagement of citizen scientists. We recommend the use of iNaturalist for future moth inventories, and as a resource for follow up meta-analyses of regional moth diversity and distributions.

## Introduction

The scaled-winged insects (Lepidoptera), which include butterflies and moths are among the most diverse order of insects, with some 160,000 named species. Some workers estimate the world Lepidopteran fauna to number as high as 350,000 species (Powell 2003). In Australia, the study of Lepidoptera has enjoyed a long history dating back to the 1800s, where natural historians and entomologists have made collections, described and compiled checklists of moth species. The most authoritative checklist of Australian Lepidoptera (Nielsen *et al*. 1996) lists 10,590 lepidoptera for Australia, although more recent estimates suggest that Australia has somewhere between 20,000 to 30,000 species of moths (Zborowski & Edwards 2007), but only about 430 butterflies (Braby 2010). Despite the huge disparity in diversity between moths and butterflies, the latter generally receives much more scientific attention and positive publicity in popular media (Franklin & Morrison 2019).

The contributions of professional lepidopterists and also naturalists have been instrumental to our understanding of regional moth diversity in the Wet Tropics Bioregion. Among this dedicated group of moth enthusiasts, the late Graeme Cocks, based in Townsville accumulated a large photographic collection of over 700 species of moths, many from the Townsville region, which have been posted on the butterflyhouse website (Herbison-Evans & Crossley 2020). The late Ian Common who wrote the first authoritative guide to moths in Australia (Common 1990) collected extensively in the region, as did entomologists Frederick Dodd, Marianne Horak, David Rentz, Bart Hacobian and others. Reaching a more popular audience, artist and lepidopterist Buck Richardson also published two guidebooks of moths in wider north Queensland region (Richardson 2008, 2016), with photographs featuring live moths rather than preserved specimen collections in biodiversity databases.

In spite of this long history of moth collections, there is still fragmentary information on the distribution of many described species, and also those of collected but not formally described species. Such a knowledge gap is known to biogeographers as the Wallacean shortfall, and is a serious impediment to effective conservation (Cardoso *et al*. 2011). For instance, although it is possible to estimate that there are 20,000-30,000 species of moths in Australia (Zborowski & Edwards 2007), it is not a straightforward matter to make an estimation on how many moths may occur in a biodiverse tropical region such as the Wet Tropics due to the Wallacean shortfall. The moth biodiversity values and conservation implications for the Wet Tropics Bioregion region is therefore yet to be properly assessed.

A related type of knowledge gap which affects biodiversity management and conservation is the Linnean shortfall, which refers to the lack of knowledge on what species exist (Cardoso *et al*. 2011). This is highly pertinent to the Australian moth fauna and particularly true for the poorly studied and so-called Microlepidoptera, an artificial grouping used by lepidopterists to describe smaller moths with wingspans <10mm (Kristensen *et al*. 2007).

Biodiversity inventories at local spatial scales can be a useful way of providing important baseline information in addressing the Wallacean shortfall and thereby helping in understanding, managing and conserving ecosystems (Dijkstra 2016). Well-documented biodiversity inventories, such as that in natural history museums (Shaffer *et al*. 1998) or digital databases (Soberón *et al*. 2007), can play crucial roles in documenting changes in species distributions. The engagement of citizens as scientists (i.e. citizen scientists) to help with biodiversity data collection has also risen in popularity, in tandem with the growing suite of different digital tools or platforms such as QuestaGame and iNaturalists, which enable citizen scientists to contribute photographic observations of wildlife and participate in generating valuable wildlife distribution data.

As a focal group, moths are highly amenable to engagement with the broader community because they are attracted to and settle around light sources at night (Zborowski & Edwards 2007), presenting themselves as easily subjects for observation and photography for people living in suburban regions close to natural habitats. Additionally, moths can also serve as important indicators of ecosystem health and change (Summerville *et al*. 2004; Highland *et al*. 2013), and therefore, accumulating long term datasets of moth seasonality and distributions can be useful for monitoring the impacts of vegetation, landuse and climate changes on the biodiversity of a locality.

The overarching aims of the study are therefore to contribute towards improving the information on the diversity and distribution of moths in the Wet Tropics bioregion of Australia (Fig. 1A), by conducting a digitally curated moth inventory of an upland rainforest site in north Queensland, whilst engaging with the broader community of citizen scientists. We pay particular attention to cases of range extensions for some species of interest in our efforts to contribute towards remedying the Wallacean shortfall of Australian moths. A secondary and minor objective of this study is also to feature images in this article and the following Part II paper (see Tng *et al*. 2020) notable moths photographed *in situ* to complement the works of Richardson (2008, 2016) and the Butterflyhouse website (Herbison-Evans & Crossley 2020).

**Figure 1.**
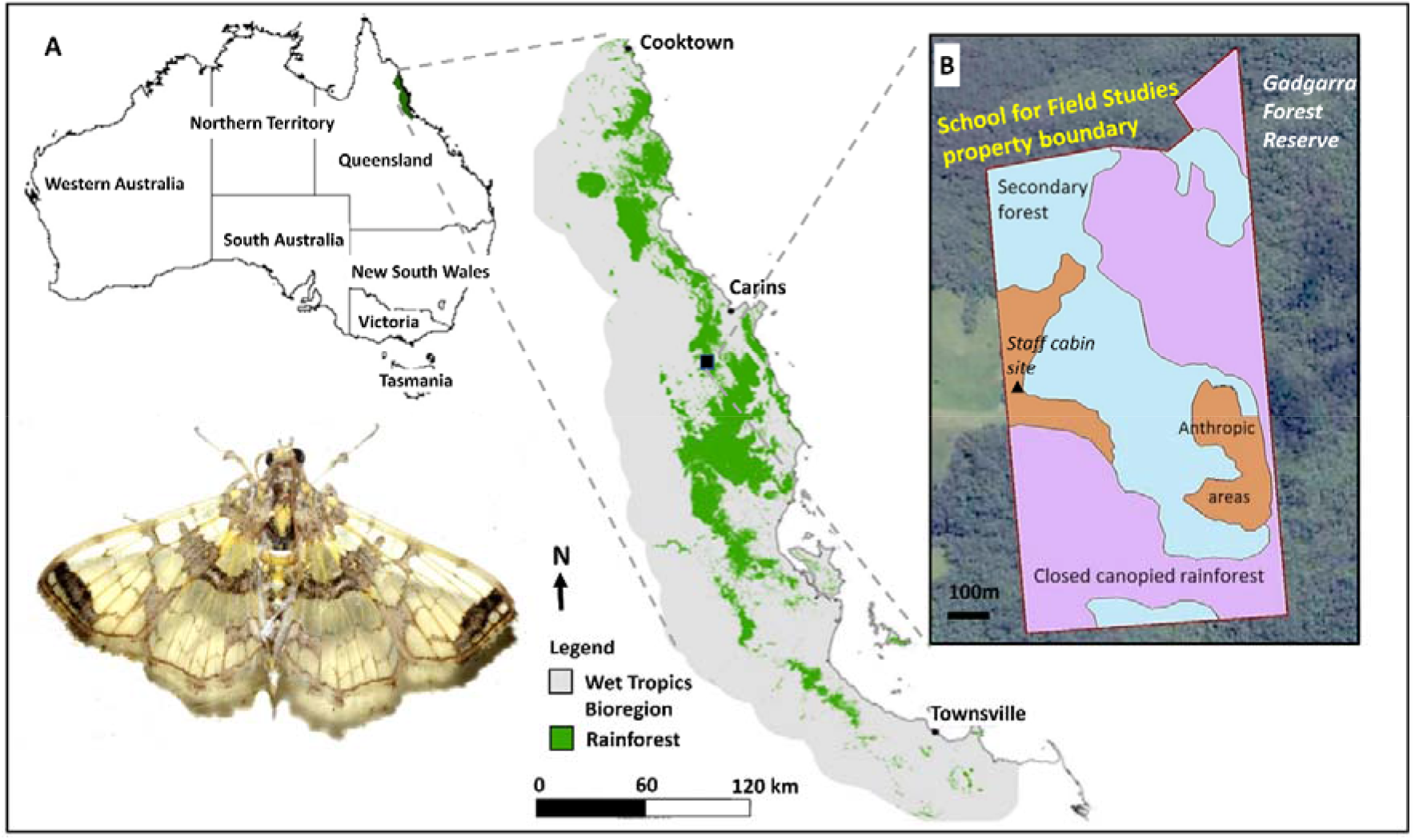
Location of the Centre for Rainforest Studies, School for Field Studies site in Gadgarra, Atherton Tablelands, Queensland where the study was conducted. (A) The general location and boundary of the Wet Tropics Bioregion, and; (B) The vegetation types in each area of the property and the location of the main staff cabin moth survey site. The featured moth is *Didymostoma aurotinctalis* (Crambidae).

## Materials and Methods

### Study Site

The Wet Tropics Bioregion of north Queensland is the geographical setting for the study (Fig. 1A), which was conducted within the corporate-owned property of the Centre for Rainforest Studies (CRS) of the School for Field Studies (17°12’S, 145°40’45”E). The CRS is located in the Atherton Tablelands region near Danbulla National Park and is approximately 13km from the nearest town Yungaburra. The site abuts the western edge of Gadgarra Forest Reserve, and consists of an area of 62 hectares of upland tropical rainforest locally known as simple mesophyll vine forest (Tracy & Webb 1975) at different stages of recovery from previous land clearing or logging activities, areas with replanted rainforest, and also a limited extent of abandoned orchard areas and built up areas (Fig. 1B; Mullin *et al*. 2020). The dominant underlying geology within the CRS is granite, with some small areas of basalt.

The main observation spot was situated at the veranda of a staff cabin that sits on a ridge and which is located near the western edge of the CRS property (Fig. 1B). The vegetation in the immediate vicinity of the staff cabin is consists of secondary forest and grassy paddocks that form part of the neighbour’s property. Also present is a thin strip of forest revegetation plantings which was planted in 2014.

### Species surveys

We conducted moth surveys on an *ad hoc* basis beginning on the 9^th^ of August 2019 until 8^th^ of August 2020. For the purpose of this work, we define a survey as an attempt to photograph all moth morphospecies observed at a sampling location, in this case, the top cabin veranda. Therefore, over the main 365 day period, we conducted moth surveys on a total of 191 nights (Fig. 2), with a gap in surveys for a few weeks in December 2019. During surveys, observations were made between 1900hrs to 2200hrs on regular forays onto the veranda to search for and to photograph moths. On a number of days where moths were abundant, these forays continued until midnight.

**Figure 2.**
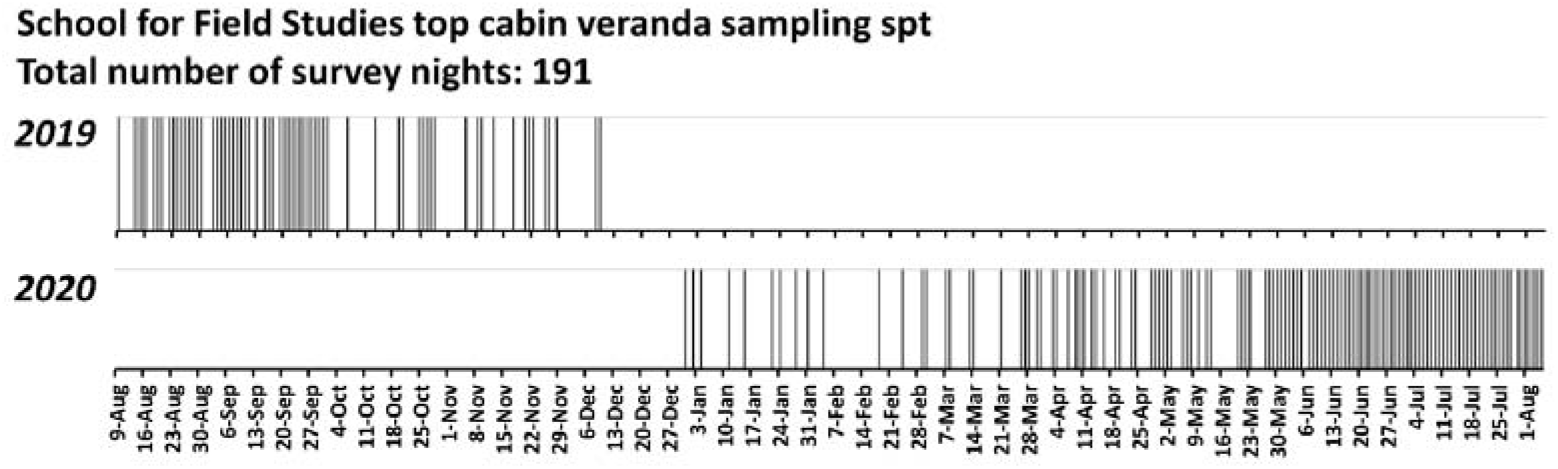
Record of days where moth surveys were conducted at the staff cabin veranda at the School for Field Studies between 2019-2020. Each thin vertical bar represents a single night’s survey.

The source of attraction for the moths was a single Phillips TLD 36 watt tube light which illuminated the 3.6m x 2.1m veranda. In keeping with the non-specialist nature of the observational study, we did not use the typical white screen method, nor did we make any physical collections of moths. Rather, we searched for and photographed moths that settled around surfaces illuminated by the light source, such as on the underside of the roofing panels, rafters, and outer walls and windows of the cabin. Moths were photographed using either a handphone camera (Samsung Galaxy S9) or a Nikon D3200 with a mounted AF Micro Nikkor 60mm lens, and most specimens were photographed to present the dorsal view of the insect, although where possible, ventral and lateral shots were taken. In most cases, fill in flash was used.

During the study period, a number of incidental moth observations were also made at other random spots within the property by the main observer (DYPT) and students who stayed at the Centre during their study abroad semester. Beyond the 8^th^ of August 2020, we continued to make incidental observations and photograph moths, with an emphasis on moth species that we had not seen during the initial one-year time frame. We report these taxa in our checklist (Supplementary Material Table S1) but we do not use them in our main analyses.

### Observation curation and identification

As part of our aim to have all the observations curated on a publicly accessible online database, all photographic observations were uploaded onto the iNaturalist platform at iNaturalist.org, either directly from the handphone app or more commonly from the computer after post-processing (see later). The platform is a joint initiative of the California Academy of Sciences and the National Geographic Society and is designed to support an online social network of people sharing biodiversity information to help each other learn about nature, or as a way of contributing to citizen science projects, institution-based initiatives, or for personal curation of biodiversity observations (Ueda 2020). Additionally, observations submitted to the iNaturalist database are shared with the Atlas of Living Australia (ALA) and the Global Biodiversity Information Facility (Seltzer 2019).

Prior to uploading observations on iNaturalist, we first processed the moth photographs in imaging software Picasa 3 to brighten or improve contrast and to crop images to fit the frame to facilitate identification. Upon uploading each photographic observation onto the platform, metadata such as date, time, and sometimes location associated with each photograph would also be automatically be reflected on the observation page. For increased accuracy, we defined the observation locations manually.

We created a project entitled “Moths at the School for Field Studies Australia” (https://www.inaturalist.org/projects/moths-at-the-school-for-field-studies-australia) into which all observations uploaded and falling within a manually-drawn polygon area specified for the site of interest (i.e. the Centre for Rainforest Studies property) would be automatically curated under that the project. For the purpose of curating observations specifically from the top cabin within the one-year sampling period, we created an addition project entitled “Veranda moth-er for a year” (https://www.inaturalist.org/projects/veranda-moth-er-for-a-year), which specified a smaller polygon encompassing only the veranda area of the top cabin and specifying the time period during which our intensive sampling occurred (9^th^ of August 2019 until 8^th^ of August 2020).

Upon uploading photographs onto the iNaturalist platform, the iNaturalist artificial intelligence interface allows observers to select suggestions of taxa that are computed to be the most likely match for the photograph observation uploaded (van Horn *et al*. 2018). Because the principal authors (DYPT and DMGA) are not lepidopteran specialists, we avoided using these machine-generated identification suggestions, but rather, entered “Lepidoptera” into the identification field. These unidentified observations would then be left to the broader community of experts, and also to coauthors VWFIII and NJF in their capacity as citizen scientists, to identify. For some observations, initial identifications were also attempted by the main observer (DYPT) by visual matching with photographs published in the literature (Common 1990; Richardson 2008, 2016) and online resources such as the Butterflyhouse website (Herbison-Evans & Crossley 2020) and the Barcode of Life Database system website (boldsystems.org; henceforth BOLD; Ratnasingham & Herbert 2007).

The iNaturalist platform lists varying levels of data quality for each observation depending on the level of consensus in identification of the observation. Regardless of whether the user who uploaded the observation has assigned a species name, observations are assigned a “Needs ID” status when they are initially uploaded. After the observation has obtained the consensus of two iNaturalist users, it becomes a “Research Grade” observation, which is suitable for biodiversity research purposes. Therefore, where only one identification was made by a user, we either solicited additional identifications from other experts, or cross checked the observation with photograph database of specimens at BOLD. If the observation was a good match for the photograph specimens in BOLD, we accepted the identification provided, thus resulting in the observation becoming “Research Grade”. The url for the taxon page at the BOLD database was then pasted into the notes field or comments section of the observation entry as a reference. Nevertheless, some observations with assigned species names still remain in a “Needs ID” status but we include this in our data analysis and species checklist because we are confident of their identification.

### Data analysis

We exported the data from the “Veranda moth-er for a year” iNaturalist project in a Microsoft Excel csv format for data analysis. The data was first put through a process of data cleaning to obtain a constrained set of observations because not all observations were suitable to be used for analysis. In cases where we chose to omit observations, it was typically because the images were of too poor a quality (due to faded markings, damaged wings, etc) to make a robust identification. In some other cases we could not ascertain if an unidentified observation already belongs to the pool of identified observation, and so we took the conservative approach to omit such observations from the analysis, rather than assigning them to species or assigning an unknown “Lepidoptera” status.

To address our objective to understand moth diversity and sampling effort, we calculated the frequency of sighting for species which is the number of surveys in which a taxon or morphospecies was sighted during the 191 survey days expressed as a percentage. We plotted a rarefied morphospecies accumulation curve by the number of surveys in the PAST 3.0 (Hammer *et al*. 2001) statistical program. For both the frequency reporting and morphospecies accumulation curve, we excluded a few morphospecies that, for various reasons, we could not be certain did not belong to an already named species in our dataset.

As part of our objective to address the “Wallacean shortfall” (distribution data shortfall) on moths, we feature noteworthy finds in the results, using the full dataset from the “Moths at the School for Field Studies Australia” project, with an emphasis on species that either represent a new record for the Wet Tropics Bioregion, a range extension, or fill a distribution gap. For our distribution analysis, we examined records of each species in the distribution map for the species in iNaturalist, the Atlas of Living Australia and in BOLD, primarily to determine if the species has ever been documented in the Atherton Tablelands or Wet Tropics region.

## Results

During the 191 nights of observations at the veranda of the staff cabin between 9^th^ Aug 2019 to 2020, we photographed and uploaded a total of 3901 observations. Among the constrained set of 3070 observations that we used for our analysis, 906 morphospecies were distinguished. Of these morphospecies, 564 were identified to species representing 477 genera and 49 moth families; 144 were identified to genus level; and 178 to higher taxonomic levels (Superfamily, family, subfamily or tribe) (Table 1). Thirty-four morphospecies were left simply as “Lepidoptera” because we were unable to assign these taxa into more refined groups, and also no iNaturalist users had suggested identifications. A total of 102 iNaturalist users, including the authors, contributed identifications. Additional incidental observations comprising of 100 morphospecies (44 named species and 56 at genus or higher taxonomic levels) were documented from the property.

**Table 1.**
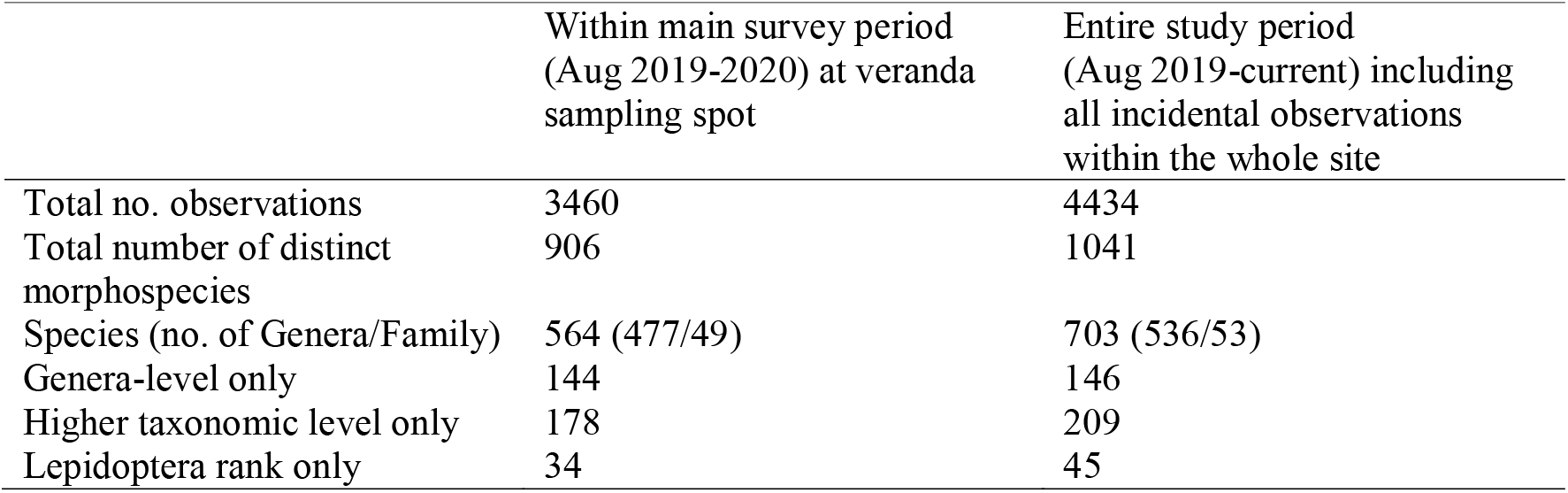
Moth species richness at the Centre for Rainforest Studies, School for Field Studies site in Gadgarra, Atherton Tablelands, Queensland.

All observations pertaining to surveys conducted at the cabin veranda within the one-year observation period (9^th^ August 2019-2020) by a single observer (DYPT) may be found at the “Veranda moth-er for a year” project page (https://www.inaturalist.org/projects/veranda-moth-er-for-a-year), while the full set of observations uploaded by multiple iNaturalist users to the current date current may be found at the “Moths for the School for Field Studies” project page (https://www.inaturalist.org/projects/moths-at-the-school-for-field-studies-australia). A complete checklist of moth morphospecies for the site and the details of digital observation curation are presented in the Supplementary Material Table S1.

### Moth frequency at the cabin veranda sampling spot

Our morphospecies accumulation curve analysis (Fig. 3A) show that we have yet to plateau in the number of species that visit the location and are still require more surveys to get a more robust estimate of the species richness of the locality. Indeed, even after the one-year survey period, we continued to observe new morphospecies. Of note also, the huge majority (497) of the morphospecies were observed only once (Fig. 3B).

**Figure 3.**
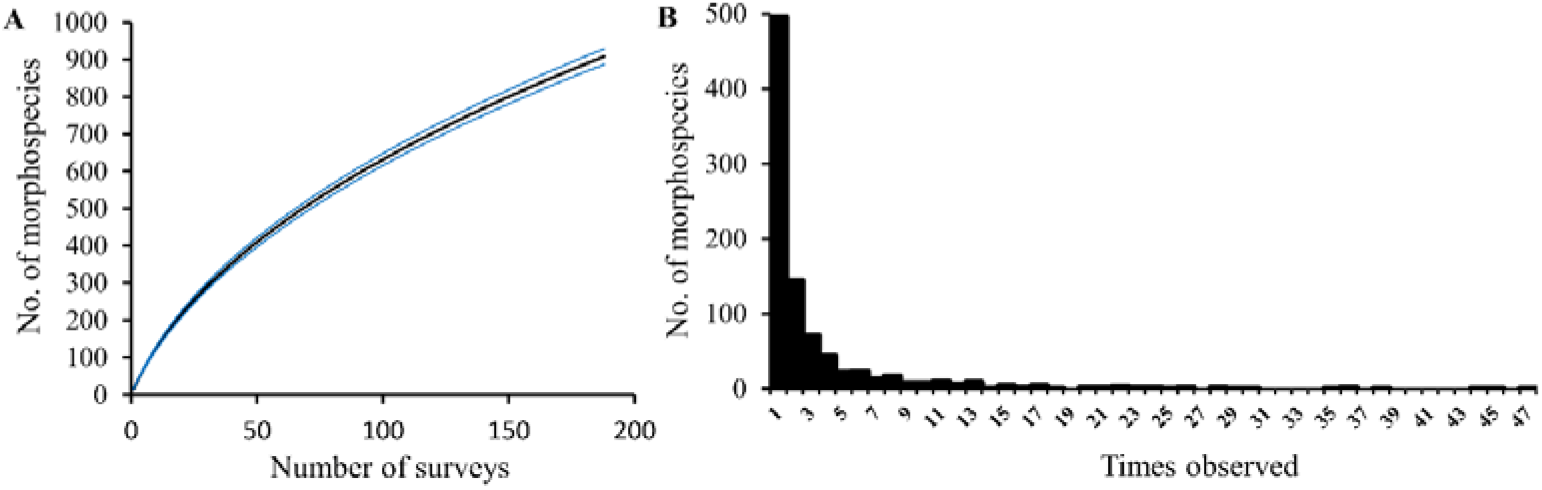
Accumulation curve for the number of mophospecies detected with number of (A) survey days and (B) frequency of species observations. Only 909 distinct species/morphospecies for which we were confident are distinct taxonomic entities were used for this analysis. The blue lines in (A) depict the standard errors of the rarefaction analysis.

The top five most frequently encountered species in a descending order of frequency are *Symmetrodes sciocosma, Endotricha mesenterialis, Mocis frugalis, Pisara hyalospila* and *Cnaphalocrocis poeyalis*. Members of the Erebidae (206 morphospecies), Geometridae (124), Crambidae (117), Noctuidae (68) and Pyralidae (61) were the most frequent visitors during the main one-year survey period.

### Noteworthy observations

Among the species observed throughout the entire observation period (Aug 2019 – Nov 2021), *Perixera* sp. AAI2525, may represent a new record for Australia (Fig. 4A), and was previously collected only from Papua New Guinea. A further nine species represented a range extension of south Queensland species (see also Tng *et al*. 2020, this issue). Some notable examples include *Callithauma pyrites*, *Conoeca guildingi*, *Dysallacta* sp. ANIC 1, *Elaphromorpha glycymilicha, Euthrausta phoenicea, Scoparia chiasta*, and *Tachystola thiasotis* (Fig. 4B-F respectively).

**Figure 4.**
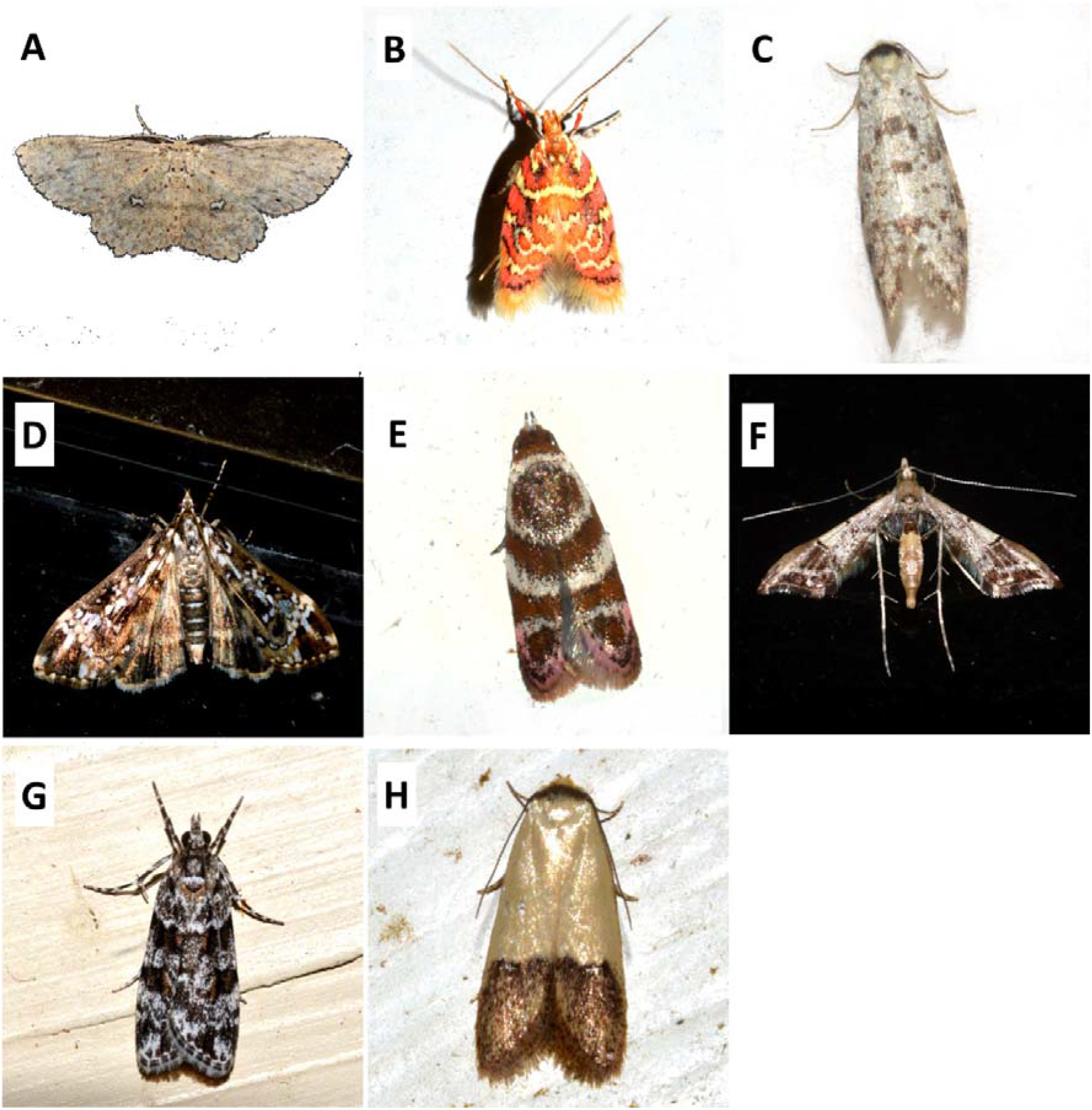
Selected moth species documented at the Centre for Rainforest Studies, School for Field Studies site in Gadgarra, Atherton Tablelands, Queensland that represent range extensions. These include a potentially new record for Australia - (A) *Perixera* sp. AAI2525 (Geometridae) and various species (B-E) that are new records for the Wet Tropics Bioregion - (B) *Callithauma pyrites* (Oecophoridae); (C) *Conoeca guildingi* (Psychidae); (D) *Dysallacta* sp. ANIC 1. (Crambidae); (E) *Elaphromorpha glycymilicha* (Oecophoridae); (F) *Euthrausta phoenicea* (Tineodidae); (G) *Scoparia chiasta* (Crambidae), and; (H) *Tachystola thiasotis* (Oecophoridae). Photographs by David Tng.

We also observed a number of species for which there are very few collections in BOLD systems or in the Atlas of Living Australia (Fig. 5). Some of these species such as *Acorotricha crystantha* (Fig. 5A) and *P. zoonophora* (Fig. 5I) may be endemic to the Wet Tropics Bioregion (Table S1). Of note, *Calamotropha* sp. 1 (Fig. 5B) and *Parapadna zoonophora* (Fig. 5I) are represented by single collections, and *Metaphoenia rhodia* (Fig. 5E) by two collections in the north Queensland region in BOLD. Others such as *Crocanthes halurga* (Fig. 5B) and *Gymnoscelis callichlora* (Fig. 5C) have distributions extending down to South Queensland, while *Nemophora panaeola* (Fig. 5F) has a highly disjunct distribution with a record from New South Wales.

**Figure 5.**
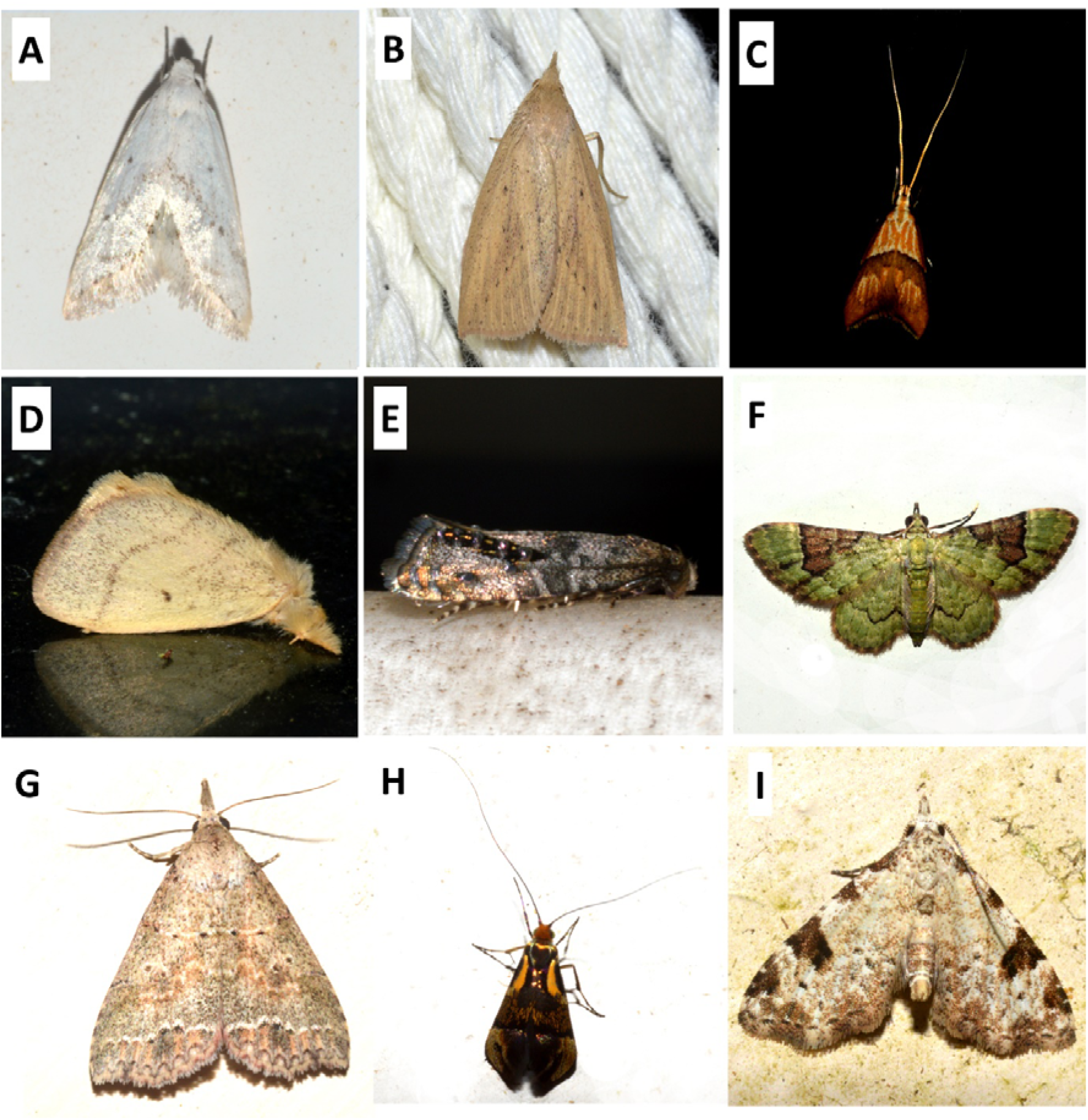
Species documented at the Centre for Rainforest Studies, School for Field Studies site in Gadgarra, Atherton Tablelands, Queensland that are known from very few records in BOLD systems and the Atlas of Living Australia. (A) *Acorotricha crystantha* (Oecophoridae); (B) *Calamotropha* sp. 1 (Crambidae); (C) *Crocanthes halurga* (Lecithoceridae) (D) *Euproctis aganopa* (Lymantriidae); (E) *Glyphipterix marmaropa* (Glyphiterigidae); (F) *Gymnoscelis callichlora* (Geometridae); (G) *Metaphoenia rhodias* (Erebidae); (H) *Nemophora panaeola* (Adelidae) and; (I) *Parapadna zoonophora*.(Erebidae). Photographs by David Tng.

Some non-native moths such as *Lantanophaga pusillidactylus* (Pterophoridae) and *Neurostrota gunniella* (Gracillariidae) were also observed during the study

### Featured moths

During the study, many moths were photographed for the first time as living specimens, rather than as pinned specimens in collections. Previously, many of these species are only featured on the BOLD systems website as photo-databased pinned specimens. Many of the moth photographs that we uploaded onto the iNaturalist platform also represent the first photograph for the species uploaded onto the system. All photographs may be accessed on the Moths for the School for Field Studies project page (https://www.inaturalist.org/projects/moths-at-the-school-for-field-studies-australia).

Unlike most moth diversity studies which exclude the Microlepidoptera, that is small moths with windspans < 1cm, we photographed numerous taxa of small moths (Fig. 6) representing at least 14 different families. These included members from the: Argyresthiidae (Fig. 16); Batrachedridae (Fig. 6B); Blastobasidae (Fig. 6C); Bucculatricidae (Fig. 6D); Choreutidae (Fig. 6E); Cosmopterigidae (Fig. 6F-G); Gelechiidae (Fig. 6I-K); Gracillariidae (Fig. 6L-M); Micropterigidae (Fig. 6N); Nepticulidae (Fig. 6O); Opostegidae (Fig. 6P); Stathmopodidae (Fig. 6Q); Tineidae (Fig. 6R-S), and; Yponomeutidae (Fig. 6T).

**Figure 6.**
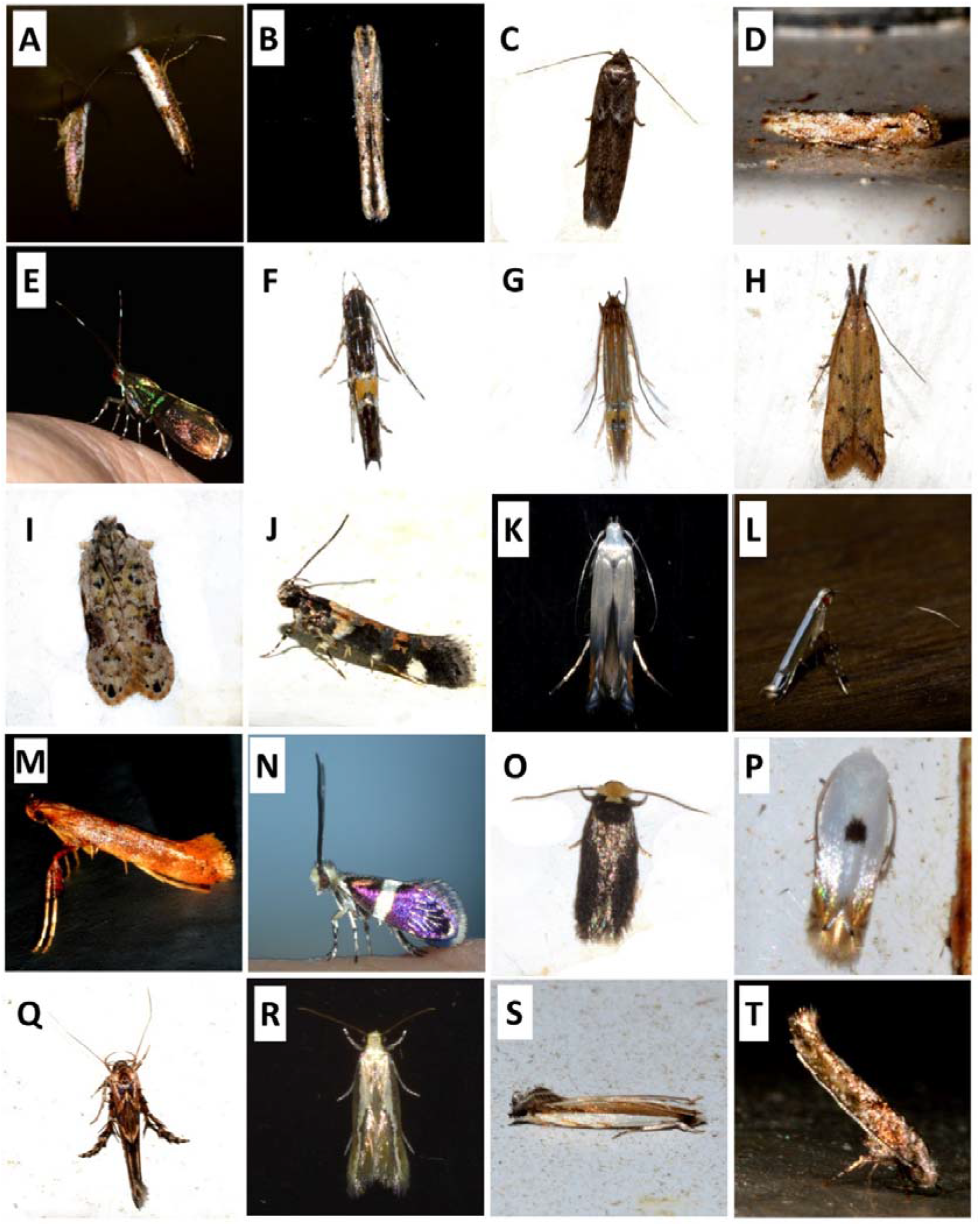
A plethora of “small moth” (Microlepidoptera) species documented at the Centre for Rainforest Studies, School for Field Studies site in Gadgarra, Atherton Tablelands, Queensland. (A) *Argyresthia notoleuca* (Argyresthiidae); (B) *Batrachedra* sp. (Batrachedridae) (C) *Blastobasis* sp. (Blastobasidae); (D) *Bucculatrix* sp. (Bucculatricidae); (E) *Saptha libanota* (Choreutidae); (F) *Cosmopterix* sp. and (G) *Labdia leucombra* (Cosmopterigidae); (H) *Dichomeris acuminata*, (I) *Hypatima* aff. *deviella*, (J) *Stegasta variana* and (K) *Thiotricha atractodes* (Gelechiidae); (L) *Acrocercops* sp. and (M) *Caloptilia* sp. (Gracillariidae); (N) *Tasmantrix thula* (Micropterigidae); (O) Unidentified sp. (Nepticulidae); (P) *Opostega* sp. (Opostegidae); (Q) *Stathmopoda castanodes* (Stathmopodidae); (R) *Opogona stenocraspeda* and (S) *Erechtias* sp. (Tineidae); (T) Unidentified sp. (Yponomeutidae).

However, many of these observations may not be reliably identified based simply on visual characters from photographs. Members of the Blastobasidae, Bucculatricidae and Nepticulidae that we observed often did not exceed 5mm in body length, and may require microscopic examination to enable further identification. Nevertheless, because the Microlepidoptera are so seldom included in moth diversity studies, the inclusion of Microlepidoptera observations in this study should form a good basis for more detailed future studies. For instance, we found a number of possibly undescribed specimens from the Gracillariidae (e.g. Fig 6M) and Opostegidae (e.g. Fig. 6P) which we could not match to existing specimens in the BOLD, and which are worthy of further taxonomic investigation.

We also feature a number of species that are not featured in Richardson’s (2008, 2016) work, nor in the extensive photographic archive of the Butterflyhouse website (Fig. 7).

**Figure 7.**
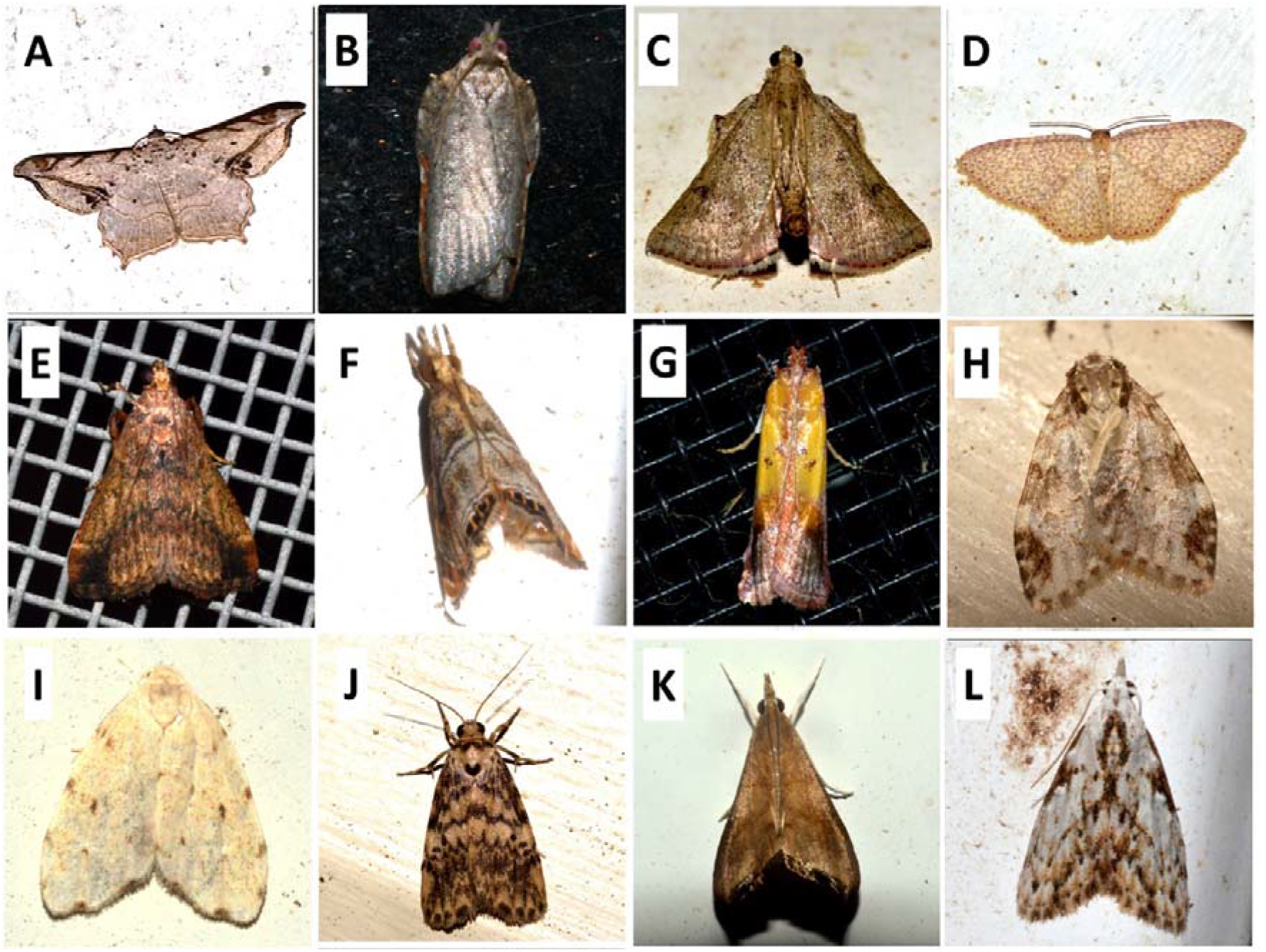
Species that are not well represented as live specimens in photographic records but documented at the Centre for Rainforest Studies, School for Field Studies site in Gadgarra, Atherton Tablelands, Queensland. (A) *Chiasmia tessellata* (Geometridae); (B) *Coeloptera gyrobathra* (Tortricidae); (C) *Endotricha lobibasalis* (Pyralidae) (D) *Eois cymatodes* (Geometridae); (E) *Gauna flavibasalis* (Pyralidae); (F) *Glaucocharis ochracealis* (Crambidae); (G) *Guastica auropurpurella* (Pyralidae); (H) *Halone camptopleura* (Erebidae); (I) *Halone ebaea* (Erebidae); (J) *Hectobrocha subnigra* (Erebidae); (K) *Hemiscopis suffusalis* (Crambidae) and; (L) *Nola sphaerospila* (Nolidae). Photographs by David Tng.

Among the 144 moth (morpho)species that we could identify to genus level, 20 could be matched to provisionally code-named species in BOLD systems (a selection featured in Fig. 8), which are based on photographed specimens curated at the Australian National Insect Collection (ANIC). Some of these species, such as *Adoxophyes* sp. D (Fig. 8A), *Araeopteron* sp. ANIC 2 (Fig. 8B) and *Progonia* sp. EF02 (Fig. 8J) were relatively frequent visitors and observed more than thrice at the Centre, but others such as *Chrysocraspedia* sp. ANIC 5 (Fig. 8C), *Corgatha* sp. ANIC 8 (Fig. 8D), and *Decticryptis* sp. ANIC 3 (Fig. 8E) were observed only once, and are also represented in collections in the BOLD systems database.

**Figure 8.**
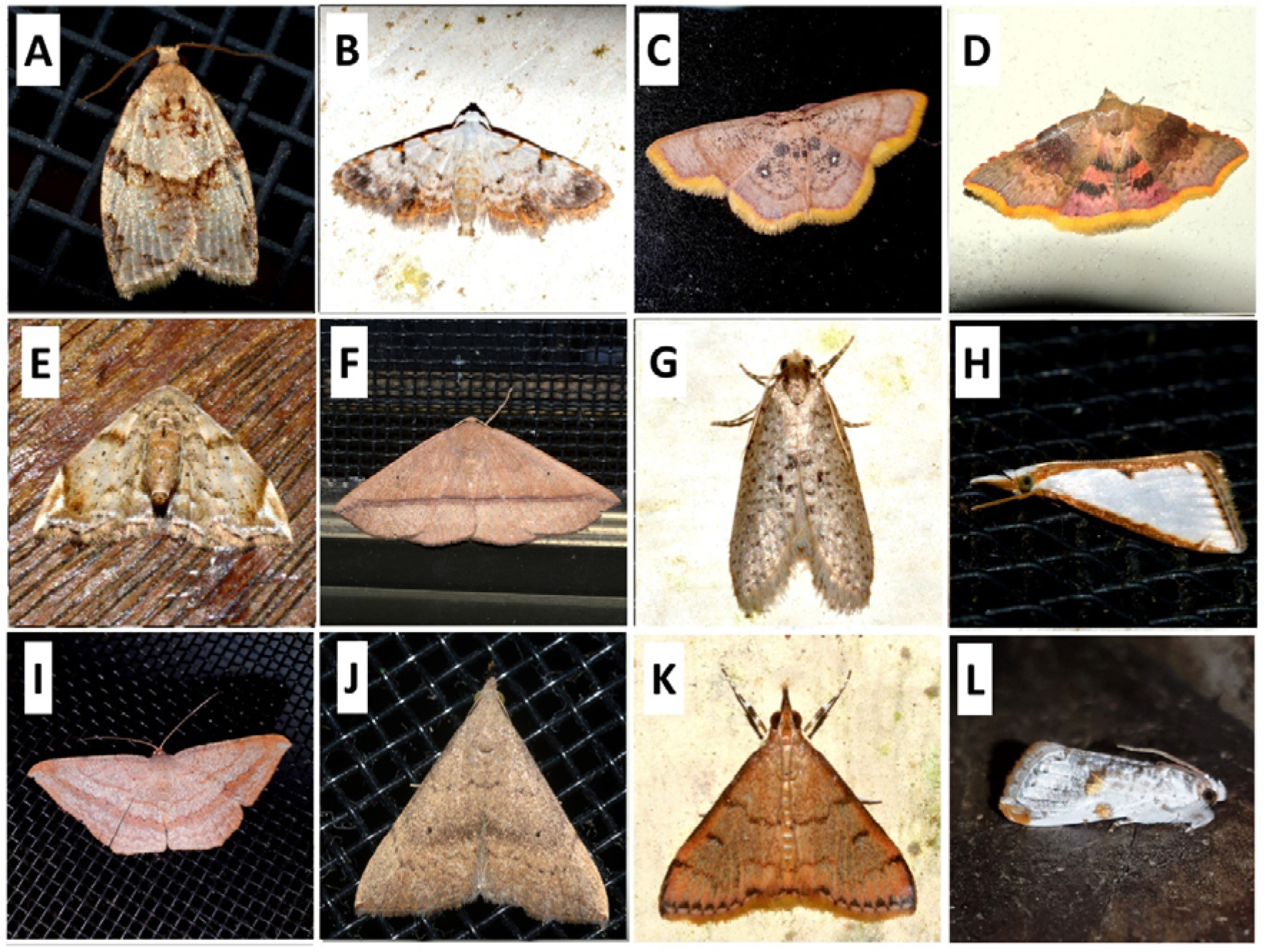
A selection of undescribed species with collections in the BOLD systems database and which are documented at the Centre for Rainforest Studies, School for Field Studies site in Gadgarra, Atherton Tablelands, Queensland. (A) *Adoxophyes* sp. D (Tortricidae); (B) *Araeopteron* sp. ANIC 2 (Erebidae); (C) *Chrysocraspedia* sp. ANIC 5 (Geometridae) (D) *Corgatha* sp. ANIC 8 (Erebidae); (E) *Decticryptis* sp. ANIC 3 (Noctuidae); (F) *Idiodes* sp. ANIC 2 (Geometridae); (G) *Lepidoscia* sp. 10 (Psychidae); (H) *Neargyria* sp. 1 (Crambidae); (I) *Probithia* sp. ANIC 1 (Geometridae); (J) *Progonia* sp. EF02 (Nolidae); (K) *Trigonoorda* sp. ANIC 4 and; (L) *Trymalitis* sp. ANIC 1 (Tortricidae). Photographs by David Tng.

## Discussion

Moth diversity studies are rare in Australia, presenting various shortfalls in knowledge that hampers our understanding of their biogeography, and impedes their conservation. In particular, studies consisting of repeated observations in a single locality are lacking and could, over the long term help reveal patterns in the seasonality of moth occurrence. Using purely photographic records identified by community experts and lodged on an online platform, we present a model for observational moth study that can add to biodiversity distribution data and engage citizen scientist participation.

### Moth species richness and occurrence frequencies

The rainforest locality where the Centre for Rainforest Studies property is situated contains a high moth species richness, as reflected by the observation of 911 distinct moth entities (563 formally named species) in the one-year sampling period at a single sampling spot and the additional 100 species documented as informal observations within the property (Table 1). In spite of this, we freely acknowledge that this moth inventory at the Centre for Rainforest Studies is an ongoing endeavour, as identifications to observations continue to be made and observations of hitherto undocumented species continue to be uploaded. It is possible that some groups of moths remain undersampled because of the observational method we used in lieu of formal collecting. We noticed for example that some moths visited the veranda but did not settle long enough on surfaces around the light source for us to photograph.

Although it is not possible to make direct comparisons between this study with published studies based on more rigorous scientific collection methods such as the use of Pennsylvania light traps (Frost 1957), a brief discussion of the moth collecting effort in north Queensland can be insightful. Indeed, one of the largest study on moth diversity in the region was a comparison between mountain ranges that included Mt Lewis in the north Queensland region (Ashton *et al*. 2016). The authors sampled rainforest canopy and understorey moths along an altitudinal gradient on Mt Lewis over six nights in total and although they excluded moths with wingspans < 1cm, they documented 17,258 individuals represented by 1134 moth species (Ashton *et al*. 2016). Additionally, they reported that moth species richness peaked at around 600m (Ashton *et al*. 2016). In another study conducted in intact rainforest and regrowth rainforest areas in the vicinity of Atherton, Kitching *et al*. (2000) collected 15,452 specimens comprising 835 species of Macrolepidoptera from three weeks of sampling each in the dry and wet seasons between 1996 to 1997, using both Pennsylvania light traps and light sheeting sampling methods.

Although we did not attempt to estimate moth abundances during our surveys, the frequency of occurrences of some moth species is worthy of mention. Two of the most frequently observed moths, *Symmetrodes sciocosma* and *Endotricha mesenterialis*, were observed on 63 and 44 nights respectively out of the 191 survey nights, and frequently with more than a single individual present (DYPT: pers. obs.). Interestingly, very little is known about the biology and food plants of these two species. The caterpillars of the third most frequently encountered moth, *Pisara hyalospila* feeds on *Elaeocarpus grandis* (Elaeocarpaceae) (Herbison-Evans & Crossley 2020), of which a number of established trees are present in the vicinity of the CRS top cabin observation site. The fourth and fifth most frequently observed moths, *Mocis frugalis* and *Cnaphalocrocis poeyalis*, are well known agricultural crop pests and their frequent presence could be due to the proximity of agricultural pastures west of the observation site.

### Addressing the “Wallacean” shortfall

Contributing towards addressing the lack of distribution data (the “Wallacean” shortfall) was a key objective of this inventory, and even though we have provided distribution data of the moth fauna for only one locality, the data represents a contribution to addressing this shortfall. Additionally, all “research grade” observations are automatically included in a larger “Moths of Queensland” iNaturalist project (https://www.inaturalist.org/projects/moths-of-queensland-australia) administered by author VWFIII, which is a resource for appreciating the distribution and seasonality of the moth fauna of the state of Queensland.

The finding of species that represent new records for the Wet Tropics bioregion and the huge number of species that we were unable to assign to any specimen in BOLD reflects the lacunae of work that still remains to be done on moth diversity and distribution in the region. Many observations may also represent a modest range extension northwards, with known collections for the Bioregion made by the late Graeme Cocks from Townsville around 250km due southeast. The previous sampling effort in Atherton by Kitching *et al*. (2000) and the checklist they published (Orr and Kitching 1999) represents the most relevant work to this study, and a cursory look revealed many overlaps in the species of the Macrolepidoptera we observed and those in their checklist. However, there were also numerous species that we observed that were missing from their checklist, reflecting some differences in species composition across localities. One potential explanation for this difference could relate to the difference in forest type as their sampling were conducted in basaltic areas which support a more structurally complex type of forest (locally known as complex notophyll vine forest). Admittedly, Orr and Kitching’s (1999) checklist also contained numerous morphospecies identified to genus or with tentative identifications, which we are unable to reconcile with our list without comparing specimens. These challenges highlight the “Linnaean” shortfalls (i.e. that many species are still not formally named) that exacerbate the moth “Wallacean” shortfall in the region.

One impediment we have observed whilst analysing distribution data is the lack of standardized information across databases. BOLD and the Atlas of Living Australia are authoritative resources to look for georeferenced distribution data for moths, but unfortunately there is still a lack of integration between the Atlas of Living Australia and BOLD, with the latter often having more complete distributional information for many of the species when we searched for information on both engines. Additionally, while for the most part an authoritative source, BOLD also suffers from some problems with data quality where sometimes photographs in the database do not belong to the species listed. It is also prudent to note also that there are still significant gaps in occurrence data in iNaturalist itself. While we did not quantify this, many of observations we uploaded represented the first observation of the species uploaded onto the iNaturalist platform although the species has been recorded in the region in BOLD and in the Atlas of Living Australia. These disparities between platforms will hopefully be addressed in the future but for now it is encouraging that Research grade observations in iNaturalist are lodged in the Atlas of Living Australia.

### Utility of the iNaturalist platform for moth inventories

One of the distinguishing features of this work is the use of the iNaturalist platform to curate photographic observations. This departs radically from other moth inventories where specimens are typically caught, preserved, and vouchered (or not). However, online curation comes with the advantage of accountability because every observation is publicly viewable, and also because identifications are vetted by a broader community of experts. Moreover, updates to the project or corrections to identifications can be made in real time, and published work related to the project benefits from wider dissemination to the public. Other side benefits of digital curation of photographed live moths is that they do away with the need to kill moths but can also complement the identification process because pinned or preserved specimens may fade or change colour with time. Admittedly, there are limits in basing identifications on purely photographic observations, but depending on study objectives, a hybrid approach involving mainly photographic vouchers and some collected specimens could be used.

In studies that rely on locality data, inaccurate georeferencing of sampling locations can negatively impact data quality. Therefore, another major advantage of using iNaturalist is that all observations can be automatically or manually georeferenced with a high level of accuracy. Such data then becomes valuable for future meta-analyses of species distributions and contributes to major efforts to digitize natural history collections to facilitate data availability online for retrieval and analysis (Beaman & Cellinese 2012). Data retrieved for meta-analyses purposes should also be put through a cleaning process to omit records of dubious quality.

The use of the iNaturalist platform for site biodiversity inventories can also be built into teaching activities and to engage students to upload wildlife or nature observations as part of academic assignments (Chamberlain *et al*. 2014). This can produce more comprehensive biodiversity inventories simply as a consequence of having more pairs of eyes looking and the ability of cover an area of interest or a wider sampling area more thoroughly. Indeed, a number of species which were not recorded at the cabin veranda sampling spot were photographed and uploaded by students who visited the Centre to undertake field courses. As a word of caution, while the potential for using iNaturalist to generated and curate biodiversity data is huge, depending on study objectives, it may be important minimize haphazard sampling by making multiple biodiversity observations over time at a given spot or spots (Callaghan *et al*. 2019), such as we have presented here. When facilitating groups of people to collect data, we also recommend that measures such as proper briefing and instruction need to be taken to ensure or improve data quality of observations from multiple users.

Finally, the utility of the iNaturalist platform for curating specimens and as a teaching tool will continue to improve, particularly because the platform is inbuilt with image recognition and machine learning technology (Gee 2017). The platform therefore “learns” to provide a more accurate suggestion of species identity with every photographic observation of a given species that is uploaded and verified. Ultimately, the utility of the platform improves with the number of people who contribute good quality observations, and therefore, using this study as a model, we thereby advocate using iNaturalist for future regional moth inventories and for long term moth distribution monitoring.

## Acknowledgements

No specific permissions were required for this study because no physical collections were made. The study also did not involve endangered or protected species. We acknowledge and give thanks to the elders of the traditional landowners past and present of the Danbulla locality. Our gratitude also goes to the various community experts, particularly Ian McMaster and Bevan Buirchell who helped us with numerous moth identification, and also staff and students of the 2019 “Fall” semester and 2020 “Spring” semesters (Alison, Bill, Sophie, Matt, Kayla, Julia, Michael and Sophie) who supplied complimentary moth observations for other spots within the CRS property.

## Supplemetary Material

Table S1. Moths recorded at the Centre for Rainforest Studies, School for Field Studies site in Gadgarra, Atherton Tablelands, Queensland.

## References

Ashton LA, Odell EH, Burwell, CJ, Maunsell SC, Nakamura A, McDonald WJF, Kitching R L. 2016. Altitudinal patterns of moth diversity in tropical and subtropical Australian rainforests. Austral Ecology 41: 197–208.

Beaman RS, Cellinese N. 2012. Mass digitization of scientific collections: New opportunities to transform the use of biological specimens and underwrite biodiversity science. ZooKeys 209: 7–17.

Braby MF. 2010. The merging of taxonomy and conservation biology: a synthesis of Australian butterfly systematics (Lepidoptera: Hesperioidea and Papilionoidea) for the 21st century. Zootaxa, 2707: 1–76.

Butler L, Kondo V, Barrows EM, Townsend EC. 1999. Effects of weather conditions and trap types on sampling for richness and abundance of forest macrolepidoptera. Environmental Entomology 28: 795–811.

Callaghan CT, Rowley JJ, Cornwell WK, Poore AG, Major RE. 2019. Improving big citizen science data: moving beyond haphazard sampling. PLoS biology 17: e3000357.

Cardoso P, Erwin TL, Borges PA, New TR. 2011. The seven impediments in invertebrate conservation and how to overcome them. Biological Conservation 144: 2647–2655.

Chamberlain A, Paxton M, Glover K, Flintham M, Price D, Greenhalgh C,… Gower A. 2014. Understanding mass participatory pervasive computing systems for environmental campaigns. Personal and Ubiquitous Computing 18: 1775–1792.

Chen IC, Shiu HJ, Benedick S, Holloway JD, Chey VK, Barlow HS,… Thomas CD. 2009. Elevation increases in moth assemblages over 42 years on a tropical mountain. Proceedings of the National Academy of Sciences 106: 1479–1483.

Common IFB. 1990. Moths of Australia. Melbourne University press: Victoria.

Dijkstra K-DB. 2016. Natural history: Restore our sense of species. Nature 533: 172–174.

Franklin DC, Morrison SC. 2019. A preliminary bioinventory of the butterflies of Talaroo Station in north Queensland’s Einasleigh Uplands. North Queensland Naturalist 49: 38–46.

Frost SW. 1957. The Pennsylvania insect light trap. Journal of Economic Entomology 50: 287–292.

Gee S. 2017. iNaturalist Launches Deep Learning-Based Identification App. https://www.i-programmer.info/news/105-artificial-intelligence/10848-inaturalist.html, viewed 31 Aug. 2020.

Hammer Ø, Harper DA, Ryan PD. 2001. PAST: paleontological statistics software package for education and data analysis. Palaeontologia Electronica 4: 9.

Herbison-Evans D, Crossley S. 2020. Australian Caterpillars and their Butterflies and Moths. http://lepidoptera.butterflyhouse.com.au, viewed 31 Aug. 2020.

Highland A, Miller JC, Jones JA. 2013. Determinants of moth diversity and community in a temperate mountain landscape: vegetation, topography, and seasonality. Ecosphere 4:129.

Kristensen NP, Scoble MJ, Karsholt OLE. 2007. Lepidoptera phylogeny and systematics: the state of inventorying moth and butterfly diversity. Zootaxa 1668: 699–747.

Mullin MV, Salecki EF, Li CR, Avinger AM, Landen EW, Thurnham JS, Pohlman CL, Tng DYP. 2020. Plant biodiversity differences between rainforest plots in different stages of recovery in the uplands of the Wet Tropics. North Queensland Naturalist 50: 25–37.

Nielsen ES, Edward DE, Tiao VR. 1995. Checklist of the Lepidoptera of Australia. East Melbourne, VIC, CSIRO Information Services.

Orr AG, Kitching RL. 1999. A checklist of Macrolepidoptera collected from rainforest and former forest areas on basalt soils on the Atherton Tableland. Australian Entomologist 26: 15–27.

Powell JA. 2003. Lepidoptera (moths, butterflies). In: Encyclopedia of Insects, eds. VH Resh, RT Carde, pp. 631–664. Academic Press: San Diego.

Ratnasingham S, Hebert P.D. 2007. BOLD: The Barcode of Life Data System (http://www.barcodinglife.org). Molecular Ecology Notes 7: 355–364.

Richardson B. 2008. Mothology: Discover the Magic – Moths and Moth Art from the World Heritage listed Wet Tropics of Queensland Australia. LeapFrogOz: Kuranda.

Richardson B. 2016. Tropical Queensland Wildlife from Dusk to Dawn Science and Art. LeapFrogOz: Kuranda.

Seltzer C. 2019. Welcome, iNaturalist Australia! https://www.inaturalist.org/blog/27851-welcome-inaturalist-australia, viewed 27 Aug. 2020.

Shaffer HB, Fisher RN, Davidson C. 1998. The role of natural history collections in documenting species declines. Trends in Ecology & Evolution 13: 27–30.

Soberón J, Jiménez R, Golubov J, Koleff P. 2007. Assessing completeness of biodiversity databases at different spatial scales. Ecography 30: 152–160.

Summerville KS, Ritter LM, Crist TO. 2004. Forest moth taxa as indicators of lepidopteran richness and habitat disturbance: a preliminary assessment. Biological Conservation 116: 9–18.

Tracy JG, Webb LJ. 1975. Vegetation of the Humid Tropical Region of North Queensland (15 maps at 1:100,000 scale + key). CSIRO Long Pocket Laboratory: Indooroopilly: Brisbane.

Ueda K. 2020. iNaturalist Research-grade Observations. iNaturalist.org. Occurrence dataset https://doi.org/10.15468/ab3s5x, viewed via GBIF.org on 26 Aug.2020.

Van Horn G, Mac Aodha O, Song Y, Cui Y, Sun C, Shepard A,… Belongie S. 2018. The inaturalist species classification and detection dataset. In Proceedings of the IEEE conference on computer vision and pattern recognition, pp. 8769–8778.

Zborowski, P, Edwards T. (Eds.). 2007. A guide to Australian moths. CSIRO PUBLISHING.

