## Supplementary Material for "Moth species richness in an upland tropical rainforest: A citizen scientist assisted study"

**The Checklist**

Species collected are listed in an alphabetical order to facilitate the utility of the list for download and for meta-analyses (Table S1). Where the prefix 'aff' is used, it indicates that the morphospecies is visually similar to a known species, or a member of a genus or family. Observations that were identified to taxonomic levels above genus are typically appended with a “gensp.” to signify that the genus and species are undetermined. In many cases, observations were matchable with specimens in BOLD which only have a provisional name. These are indicated with “ANIC”, referring to their presence in the Australian National Insect Collection or with “BOLD” indicated after sp. Where unidentified specimens were not matchable with specimens in BOLD, the specimen is given a placeholder name typically appended with a “CRS” (thus *Genus* CRS1 and so forth) or a short descriptive epithet (e.g. Lepidoptera Magneto).

We freely acknowledge that some of the moths which we were unable to identify may already be formally descripted, in which case, we lacked the expertise to provide an identification and/or no identifications from the broader iNaturalist community has been forthcoming. However, we believe also that many represent undescribed, or potentially undocumented species.

All digital photograph accessions are accessible online at the “Moths for the School for Field Studies” project page (<https://www.inaturalist.org/projects/moths-at-the-school-for-field-studies-australia>). An editable copy of the checklist is available upon request from the corresponding author (DYPT).

**Table S1. Moths recorded at the Centre for Rainforest Studies, School for Field Studies site in Gadgarra, Atherton Tablelands, Queensland.**
See main text for more details of surveys and context for range extension. The frequency of occurrence (%) of species surveyed during the main one-year survey period in Aug 2019-2020 (100*[n/191 survey nights]) are presented, otherwise observations are incidental and extended beyond Aug 2020 to Dec 2021. The Digital accession code can be used to peruse the photograph observation on iNaturalist by appending the code it to the end of the url: https://www.inaturalist.org/observations/

| **Scientific name** | **Family** | **Frequency of occurrence (%)** | **Range  extension?** | **Digital accession code** |
| --- | --- | --- | --- | --- |
| *Nemophora panaeola* | Adelidae | incidental |  | 58100711 |
| *Alucita phricodes* | Alucitidae | 0.52 |  | 56289578 |
| *Anthela astata* | Anthelidae | 1.57 |  | 54210858 |
| *Anthela deficiens* | Anthelidae | 1.05 |  | 50621740 |
| *Anthela varia* | Anthelidae | 0.52 |  | 53792887 |
| *Argyresthia notoleuca* | Argyresthiidae | 0.52 |  | 54907781 |
| *Batrachedra sp. CRS1* | Batrachedridae | 1.05 |  | 51854439 |
| *Batrachedra sp. CRS2* | Batrachedridae | 2.09 |  | 56618447 |
| *Batrachedra sp. CRS3* | Batrachedridae | 5.76 |  | 33125081 |
| *Blastobasis sp. CRS1* | Blastobasidae | 0.52 |  | 55124321 |
| *Blastobasis sp. CRS2* | Blastobasidae | 0.52 |  | 54367858 |
| *Blastobasis sp. CRS3* | Blastobasidae | 0.52 |  | 54022472 |
| *Blastobasis sp. CRS4* | Blastobasidae | 0.52 |  | 55224245 |
| *Bucculatrix sp.* | Bucculatricidae | 0.52 |  | 55695035 |
| *Saptha libanota* | Choreutidae | 1.57 |  | 47382218 |
| *Cosmopterigidae sp. CRS1* | Cosmopterigidae | incidental |  | 59314762 |
| *Cosmopterix sp.* | Cosmopterigidae | 4.71 |  | 33125054 |
| *Labdia deliciosella* | Cosmopterigidae | incidental |  | 61894728 |
| *Labdia leucombra* | Cosmopterigidae | 0.52 |  | 56327477 |
| *Labdia sp. CRS1* | Cosmopterigidae | incidental |  | 59120498 |
| *Pyroderces sp. CRS1* | Cosmopterigidae | 0.52 |  | 55466648 |
| *Endoxyla sp. CRS1* | Cossidae | 0.52 |  | 49935775 |
| *Zeuzera aeglospila* | Cossidae | 0.52 |  | 36282211 |
| *Achyra affinitalis* | Crambidae | 0.52 |  | 31780452 |
| *Achyra nigrirenalis* | Crambidae | 0.52 |  | 31605676 |
| *Aetholix flavibasalis* | Crambidae | 1.05 |  | 48898529 |
| *Agrioglypta eurytusalis* | Crambidae | 0.52 |  | 56979921 |
| *Agrioglypta excelsalis* | Crambidae | 0.52 |  | 43305150 |
| *Agrioglypta itysalis* | Crambidae | 2.62 |  | 50124326 |
| *Agrotera pictalis* | Crambidae | 0.52 |  | 46911860 |
| *Ambias eromenalis* | Crambidae | incidental |  | 100488486 |
| *Analyta albicillalis* | Crambidae | incidental |  | 100488487 |
| *Antigastra catalaunalis* | Crambidae | 1.57 |  | 48974155 |
| *Apoblepta epicharis* | Crambidae | 0.52 |  | 49085843 |
| *Archernis callixantha* | Crambidae | 0.52 |  | 51172695 |
| *Ategumia adipalis* | Crambidae | 0.52 |  | 52204581 |
| *Bradina admixtalis* | Crambidae | 12.04 |  | 56924945 |
| *Calamotropha paludella* | Crambidae | 1.05 |  | 51470195 |
| *Calamotropha sp. 1 in BOLD* | Crambidae | 1.57 |  | 49808174 |
| *Circobotys occultilinea* | Crambidae | 0.52 |  | 54907771 |
| *Cirrhochrista caconalis* | Crambidae | 1.57 |  | 48848484 |
| *Cirrhochrista sp. 2 in BOLD* | Crambidae | 0.52 |  | 53314996 |
| *Cirrhochrista sp. CRS1* | Crambidae | 2.09 |  | 55362862 |
| *Cnaphalocrocis* | Crambidae | 0.52 |  | 57013890 |
| *Cnaphalocrocis bilinealis* | Crambidae | 1.05 |  | 54221830 |
| *Cnaphalocrocis medinalis* | Crambidae | 2.09 |  | 40357007 |
| *Cnaphalocrocis poeyalis* | Crambidae | 19.9 |  | 30806631 |
| *Conogethes ersealis* | Crambidae | 5.24 |  | 32878426 |
| *Conogethes haemactalis* | Crambidae | 0.52 |  | 46476855 |
| *Conogethes semifascialis* | Crambidae | 0.52 |  | 43305171 |
| *Crambidae aff. Scoparinae CRS1* | Crambidae | 3.14 |  | 31464721 |
| *Crambidae gensp. CRS1* | Crambidae | incidental |  | 58115646 |
| *Culladia cuneiferellus* | Crambidae | 5.24 |  | 46358676 |
| *Desmia discrepans* | Crambidae | 0.52 |  | 55900019 |
| *Diaphania indica* | Crambidae | 0.52 |  | 43299666 |
| *Diasemia accalis* | Crambidae | 4.19 |  | 46360208 |
| *Diasemiopsis ramburialis* | Crambidae | 0.52 |  | 48898464 |
| *Diathrausta picata* | Crambidae | 0.52 |  | 49097030 |
| *Didymostoma aurotinctalis* | Crambidae | 1.57 |  | 56289586 |
| *Dysallacta sp. ANIC 1* | Crambidae | 0.52 | Yes | 56618438 |
| *Ebulea perflavalis* | Crambidae | 0.52 |  | 53593766 |
| *Ectadiosoma straminea* | Crambidae | incidental |  | 61944223 |
| *Euclasta gigantalis* | Crambidae | incidental |  | 57102508 |
| *Eudonia aphrodes* | Crambidae | incidental |  | 57102506 |
| *Eurrhyparodes tricoloralis* | Crambidae | 4.19 |  | 31780461 |
| *Glaucocharis molydocrossa* | Crambidae | 1.57 |  | 57239551 |
| *Glaucocharis ochracealis* | Crambidae | 0.52 |  | 50124317 |
| *Glaucocharis queenslandensis* | Crambidae | incidental |  | 59948187 |
| *Glyphodes actorionalis* | Crambidae | 0.52 |  | 56618522 |
| *Glyphodes apiospila* | Crambidae | 0.52 |  | 51558236 |
| *Glyphodes callipona* | Crambidae | 0.52 |  | 51211034 |
| *Glyphodes doleschalii* | Crambidae | 0.52 |  | 49907503 |
| *Glyphodes flavizonalis* | Crambidae | 0.52 |  | 30918323 |
| *Glyphodes stolalis* | Crambidae | 0.52 |  | 33125459 |
| *Haritalodes obliqualis* | Crambidae | incidental |  | 82091614 |
| *Heliothela ophideresana* | Crambidae | 1.05 |  | 56583728 |
| *Hemiscopis suffusalis* | Crambidae | 0.52 |  | 33125034 |
| *Hemiscopis violacea* | Crambidae | 1.05 |  | 56272217 |
| *Herpetogramma licarsisalis* | Crambidae | 18.85 |  | 56289562 |
| *Herpetogramma sp.* | Crambidae | 0.52 |  | 31844774 |
| *Hyalobathra aequalis* | Crambidae | 0.52 |  | 56583715 |
| *Hyalobathra brevialis* | Crambidae | 1.57 |  | 51558233 |
| *Hyalobathra crenulata* | Crambidae | 1.57 |  | 31844755 |
| *Hyalobathra unicolor* | Crambidae | 1.05 |  | 55377947 |
| *Hydriris chalybitis* | Crambidae | 5.76 |  | 33123754 |
| *Hydrorybina polusalis* | Crambidae | 1.05 |  | 53035507 |
| *Isocentris filalis* | Crambidae | incidental |  | 97894408 |
| *Leucinodes orbonalis* | Crambidae | 1.57 |  | 49412452 |
| *Mabra eryxalis* | Crambidae | 1.57 |  | 50243699 |
| *Margarosticha sphenotis* | Crambidae | 3.14 |  | 45858834 |
| *Maruca vitrata* | Crambidae | 4.71 |  | 50621764 |
| *Meroctena staintonii* | Crambidae | incidental |  | 97595901 |
| *Merodictya marmorata* | Crambidae | incidental |  | 59419163 |
| *Metasia sp. CRS1* | Crambidae | 0.52 |  | 56466059 |
| *Metasia spilocrossa* | Crambidae | 1.05 |  | 55900039 |
| *Metoeca foedalis* | Crambidae | 1.57 |  | 49085759 |
| *Musotiminae sp. CRS1* | Crambidae | 0.52 |  | 53433573 |
| *Nacoleia glageropa* | Crambidae | 1.05 |  | 52136086 |
| *Nacoleia rhoeoalis* | Crambidae | 15.71 |  | 30918319 |
| *Neargyria sp. 1 in BOLD* | Crambidae | 0.52 |  | 56327478 |
| *Nomophila corticalis* | Crambidae | 0.52 |  | 31152531 |
| *Omiodes basalticalis* | Crambidae | 0.52 |  | 54341116 |
| *Omiodes diemenalis* | Crambidae | 7.33 |  | 30918310 |
| *Omiodes indicata* | Crambidae | 13.61 |  | 31096116 |
| *Omiodes surrectalis* | Crambidae | 0.52 |  | 52724368 |
| *Orphanostigma* | Crambidae | 0.52 |  | 49808266 |
| *Ostrinia furnacalis* | Crambidae | 1.05 |  | 31203916 |
| *Paliga ignealis* | Crambidae | 1.05 |  | 48898559 |
| *Palpita annulata* | Crambidae | 1.05 |  | 33125049 |
| *Palpita horakae* | Crambidae | 1.05 |  | 50887228 |
| *Palpita hyaloptila* | Crambidae | incidental |  | 62842288 |
| *Palpita margaritacea* | Crambidae | 1.05 |  | 48898433 |
| *Palpita pajnii* | Crambidae | 0.52 |  | 50045244 |
| *Palpita rhodocosta* | Crambidae | 0.52 |  | 54707524 |
| *Parapoynx stagnalis* | Crambidae | 1.57 |  | 47382180 |
| *Parotis incurvata* | Crambidae | 0.52 |  | 33125015 |
| *Parotis marginata* | Crambidae | 1.57 |  | 53315024 |
| *Parotis suralis* | Crambidae | 0.52 |  | 49723577 |
| *Patania balteata* | Crambidae | 0.52 |  | 49917964 |
| *Phenacodes aleuropa* | Crambidae | 1.57 |  | 33125017 |
| *Piletocera meekii* | Crambidae | 0.52 |  | 47382212 |
| *Poliobotys ablactalis* | Crambidae | 0.52 |  | 57013860 |
| *Prophantis adusta* | Crambidae | 0.52 |  | 53315046 |
| *Prorodes mimica* | Crambidae | 1.05 |  | 30918312 |
| *Pyrausta phoenicealis* | Crambidae | 1.57 |  | 50023769 |
| *Pyraustinae gensp. CRS1* | Crambidae | 0.52 |  | 52204473 |
| *Pyraustinae gensp. CRS2* | Crambidae | 1.05 |  | 31780451 |
| *Rhimphalea lindusalis* | Crambidae | 12.57 |  | 30918321 |
| *Samea multiplicalis* | Crambidae | 0.52 |  | 53819475 |
| *Sameodes cancellalis* | Crambidae | 1.57 |  | 31780442 |
| *Sameodes iolealis* | Crambidae | 0.52 |  | 56312842 |
| *Scirpophaga nivella* | Crambidae | 0.52 |  | 53207541 |
| *Scoparia chiasta* | Crambidae | 4.19 | yes | 52345712 |
| *Spilomelinae gensp. CRS1* | Crambidae | 0.52 |  | 55465197 |
| *Spilomelinae gensp. CRS2* | Crambidae | 0.52 |  | 52435469 |
| *Spilomelinae gensp. CRS3* | Crambidae | 0.52 |  | 56023557 |
| *Spoladea recurvalis* | Crambidae | 7.85 |  | 31780476 |
| *Stemorrhages marthesiusalis* | Crambidae | incidental |  | 60080358 |
| *Strepsinoma croesusalis* | Crambidae | 0.52 |  | 52915153 |
| *Syllepte ridopalis* | Crambidae | 1.57 |  | 47382244 |
| *Talanga sexpunctalis* | Crambidae | incidental |  | 100135956 |
| *Tetrernia teminitis* | Crambidae | 6.81 |  | 50131501 |
| *Theila siennata* | Crambidae | 1.05 |  | 49808290 |
| *Trichophysetis sp. CRS1* | Crambidae | 0.52 |  | 49777870 |
| *Trichophysetis sp. CRS2* | Crambidae | 1.05 |  | 54707593 |
| *Trigonoorda sp. ANIC4* | Crambidae | 3.14 |  | 50621743 |
| *Uresiphita insulicola* | Crambidae | 0.52 |  | 56466048 |
| *Uresiphita ornithopteralis* | Crambidae | 0.52 |  | 52879174 |
| *Barantola panarista* | Depressariidae | 0.52 |  | 50771410 |
| *Barantola pulcherrima* | Depressariidae | incidental |  | 61786570 |
| *Depressariidae gensp. CRS1* | Depressariidae | 1.05 |  | 53317751 |
| *Eutorna pabulicola* | Depressariidae | 0.52 |  | 55861697 |
| *Hypsidia robinsoni* | Drepanidae | incidental |  | 59108465 |
| *Tridrepana lunulata* | Drepanidae | 1.05 |  | 50031218 |
| *Leptozestis sp. CRS1* | Elachistidae | 1.05 |  | 56295623 |
| *Epermenia commonella* | Epermeniidae | 0.52 | Yes, slight | 56728626 |
| *Acantholipes trajecta* | Erebidae | 1.05 |  | 30501901 |
| *Achaea janata* | Erebidae | 4.71 |  | 50004079 |
| *Achaea serva* | Erebidae | 1.05 |  | 48898449 |
| *Acyphas chionitis* | Erebidae | 0.52 |  | 47351481 |
| *Adrapsa ablualis* | Erebidae | 14.66 |  | 31096134 |
| *Agamana cavatalis* | Erebidae | 0.52 |  | 50012338 |
| *Agape chloropyga* | Erebidae | 0.52 |  | 53819476 |
| *Alapadna pauropis* | Erebidae | 2.09 |  | 54707535 |
| *Alophosoma hypoxantha* | Erebidae | 1.05 |  | 51349629 |
| *Alophosoma sp. CRS1* | Erebidae | 1.05 |  | 47351423 |
| *Alophosoma syngenes* | Erebidae | 3.14 |  | 49723553 |
| *Amata leucacma* | Erebidae | incidental |  | 61479957 |
| *Ameleta panochra* | Erebidae | 0.52 |  | 56618509 |
| *Amerila alberti* | Erebidae | 0.52 |  | 50874570 |
| *Anomis definata* | Erebidae | 0.52 |  | 49808200 |
| *Anticarsia irrorata* | Erebidae | 0.52 |  | 50105038 |
| *Araeopteron aff. pleurotypa* | Erebidae | 0.52 |  | 48898461 |
| *Araeopteron canescens* | Erebidae | 0.52 |  | 56618445 |
| *Araeopteron epiphracta* | Erebidae | 1.05 |  | 50045252 |
| *Araeopteron sp. ANIC 2* | Erebidae | 4.19 |  | 50105030 |
| *Araeopteron sp. ANIC 6* | Erebidae | 0.52 |  | 49085709 |
| *Araeopteron sp. CRS1* | Erebidae | 1.05 |  | 54341119 |
| *Araeopteron sp. CRS2* | Erebidae | 0.52 |  | 54886185 |
| *Araeopteron sp. CRS3* | Erebidae | 0.52 |  | 49917908 |
| *Arctornis lucens* | Erebidae | 0.52 |  | 55899999 |
| *Arctornis sp. CRS1* | Erebidae | 5.24 |  | 50550165 |
| *Argina astrea* | Erebidae | 4.19 |  | 31605661 |
| *Arthisma scissuralis* | Erebidae | 0.52 |  | 47382267 |
| *Artigisa impropria* | Erebidae | 0.52 |  | 50669056 |
| *Asota heliconia* | Erebidae | 2.62 |  | 31152535 |
| *Asota orbona* | Erebidae | 0.52 |  | 32523846 |
| *Asura semivitrea* | Erebidae | 5.76 |  | 50621723 |
| *Athyrma subpunctata* | Erebidae | 0.52 |  | 48898421 |
| *Avatha discolor* | Erebidae | 0.52 |  | 48898541 |
| *Axiocteta oenoplex* | Erebidae | 3.14 |  | 30918322 |
| *Bastilla absentimacula* | Erebidae | 1.05 |  | 53792901 |
| *Bastilla serratilinea* | Erebidae | 0.52 |  | 48898403 |
| *Bastilla solomonensis* | Erebidae | 1.05 |  | 54210856 |
| *Bocula odontosema* | Erebidae | 0.52 |  | 45858852 |
| *Boletobiinae gensp. CRS2* | Erebidae | 0.52 |  | 49097039 |
| *Britha biguttata* | Erebidae | 0.52 |  | 49307815 |
| *Brunia dorsalis* | Erebidae | 0.52 |  | 48974109 |
| *Buzara infractafinis* | Erebidae | 2.09 |  | 54521438 |
| *Buzara latizona* | Erebidae | 0.52 |  | 50247988 |
| *Calamidia hirta* | Erebidae | 2.62 |  | 33474921 |
| *Calyptra minuticornis* | Erebidae | 0.52 |  | 53819477 |
| *Cerynea trogobasis* | Erebidae | 2.62 |  | 54707515 |
| *Ceryx sp. CRS1* | Erebidae | 1.05 |  | 50202150 |
| *Ceryx sphenodes* | Erebidae | 0.52 |  | 45858851 |
| *Chalciope alcyona* | Erebidae | 2.09 |  | 31780438 |
| *Chrysomesia lophoptera* | Erebidae | 2.09 |  | 52999306 |
| *Corgatha sp ANIC 8* | Erebidae | 0.52 |  | 49924277 |
| *Creatonotos gangis* | Erebidae | 9.42 |  | 31780439 |
| *Crioa hades* | Erebidae | 1.57 |  | 53207396 |
| *Cyana meyricki* | Erebidae | 0.52 |  | 43305179 |
| *Cyana obscura* | Erebidae | 4.19 |  | 53433569 |
| *Dahlia capnobela* | Erebidae | 1.05 |  | 53016860 |
| *Damias pelochroa* | Erebidae | 0.52 |  | 56895215 |
| *Daona detersalis* | Erebidae | 1.57 |  | 56577865 |
| *Decticryptis sp. ANIC 3 in BOLD* | Erebidae | incidental |  | 59314737 |
| *Donuca castalia* | Erebidae | 0.52 |  | 33123765 |
| *Donuca rubropicta* | Erebidae | 4.71 |  | 30806627 |
| *Dura niveus* | Erebidae | incidental |  | 84542090 |
| *Dura ochrias* | Erebidae | 0.52 |  | 50271563 |
| *Dysgonia monogona* | Erebidae | 0.52 |  | 50199287 |
| *Dysgonia senex* | Erebidae | 0.52 |  | 50108636 |
| *Eilema plana* | Erebidae | 1.05 |  | 54221832 |
| *Enispa niviceps* | Erebidae | 0.52 |  | 31780440 |
| *Enispa parva* | Erebidae | 1.57 |  | 49307820 |
| *Enispa prolectus* | Erebidae | 0.52 |  | 49723552 |
| *Enispa sp. CRS1* | Erebidae | 0.52 |  | 43305180 |
| *Ercheia ekeikei* | Erebidae | 1.57 |  | 50243697 |
| *Erebidae gensp. CRS1* | Erebidae | 0.52 |  | 53314999 |
| *Erebidae gensp. CRS2* | Erebidae | 0.52 |  | 51854437 |
| *Erebidae gensp. CRS3* | Erebidae | 0.52 |  | 49825650 |
| *Erebidae gensp. CRS4* | Erebidae | incidental |  | 57095053 |
| *Erebidae gensp. CRS5* | Erebidae | incidental |  | 58480341 |
| *Erebus crepuscularis* | Erebidae | 0.52 |  | 50560432 |
| *Eressa aff. rhysoptila* | Erebidae | 0.52 |  | 46476835 |
| *Eressa megalospilia* | Erebidae | 1.05 |  | 50338638 |
| *Eressa rhysoptila* | Erebidae | 5.76 |  | 50031211 |
| *Eressa strepsimeris* | Erebidae | 0.52 |  | 49085772 |
| *Ericeia goniosema* | Erebidae | 0.52 |  | 52724337 |
| *Ericeia inangulata* | Erebidae | 2.62 |  | 50024918 |
| *Ericeia pertendens* | Erebidae | incidental |  | 59948198 |
| *Ericeia plaesiodes* | Erebidae | 1.05 |  | 48898519 |
| *Esthlodora versicolor* | Erebidae | 0.52 |  | 32878669 |
| *Eublemma abrupta* | Erebidae | 2.09 |  | 48898538 |
| *Eublemma accedens* | Erebidae | 1.57 |  | 55976833 |
| *Eublemma pectorora* | Erebidae | 0.52 |  | 55573922 |
| *Eublemma roseana* | Erebidae | 1.57 |  | 55900048 |
| *Eudocima phalonia* | Erebidae | 1.05 |  | 50031208 |
| *Eudocima salaminia* | Erebidae | 1.05 |  | 44581589 |
| *Eulocastra fasciata* | Erebidae | 0.52 |  | 48974129 |
| *Euproctis crocea* | Erebidae | 0.52 |  | 55980655 |
| *Euproctis epaxia* | Erebidae | incidental |  | 100135969 |
| *Euproctis galactopis* | Erebidae | 1.57 |  | 52999341 |
| *Euproctis holoxutha* | Erebidae | 1.05 |  | 31152542 |
| *Euproctis lucifuga* | Erebidae | 1.05 |  | 43299692 |
| *Euproctis trispila* | Erebidae | 1.05 |  | 50271553 |
| *Gesonia obeditalis* | Erebidae | 10.47 |  | 43305134 |
| *Goniosema anguliscripta* | Erebidae | 0.52 |  | 31780430 |
| *Goniosema euraphota* | Erebidae | 1.05 |  | 31464710 |
| *Gonitis involuta* | Erebidae | 2.09 |  | 33123748 |
| *Grammodes oculicola* | Erebidae | 0.52 |  | 50108627 |
| *Grammodes quaesita* | Erebidae | 1.05 |  | 43305133 |
| *Gymnasura saginaea* | Erebidae | 0.52 |  | 53792905 |
| *Halone camptopleura* | Erebidae | 0.52 |  | 33125032 |
| *Halone ebaea* | Erebidae | 0.52 |  | 49808225 |
| *Halone sejuncta* | Erebidae | 10.99 |  | 31203909 |
| *Hectobrocha subnigra* | Erebidae | 5.76 |  | 31096124 |
| *Heliosia sp. CRS1* | Erebidae | 0.52 |  | 56289633 |
| *Hemonia simillima* | Erebidae | incidental |  | 58115641 |
| *Herminiinae aff Nodaria sp CRS1* | Erebidae | 1.05 |  | 53016856 |
| *Hesychopa chionora* | Erebidae | incidental |  | 60314408 |
| *Heterallactis stenochrysa* | Erebidae | 2.09 |  | 52927302 |
| *Hulodes caranea* | Erebidae | 1.05 |  | 36497655 |
| *Hydrillodes dimissalis* | Erebidae | 0.52 |  | 47382295 |
| *Hydrillodes funestalis* | Erebidae | 5.24 |  | 49935772 |
| *Hypena conscitalis* | Erebidae | 10.47 |  | 56370342 |
| *Hypena gonospilalis* | Erebidae | incidental |  | 60024755 |
| *Hypena masurialis* | Erebidae | 6.28 |  | 31096132 |
| *Hyposada hydrocampata* | Erebidae | 1.57 |  | 49097021 |
| *Ischyja albata* | Erebidae | incidental |  | 41251267 |
| *Ischyja manlia* | Erebidae | 0.52 |  | 45858857 |
| *Laelia obsoleta* | Erebidae | 0.52 |  | 43299695 |
| *Lambula pristina* | Erebidae | 0.52 |  | 54707549 |
| *Lambula sp. Darkmarked* | Erebidae | 0.52 |  | 54907767 |
| *Lambula sp. Darkplain* | Erebidae | 2.62 |  | 52204488 |
| *Lambula transcripta* | Erebidae | 7.85 |  | 53246579 |
| *Laspeyria poecilota* | Erebidae | 3.14 |  | 47351397 |
| *Lemyra maculifascia* | Erebidae | 1.05 |  | 49407713 |
| *Lithosiini aff. Ateucheta* | Erebidae | 1.05 |  | 54694830 |
| *Lithosiini aff. Eilema* | Erebidae | 12.04 |  | 33123735 |
| *Lithosiini aff. Goniosema* | Erebidae | 0.52 |  | 49720670 |
| *Lithosiini aff. Meteura* | Erebidae | 0.52 |  | 56370388 |
| *Lithosiini aff. Schistophleps* | Erebidae | 1.57 |  | 55124345 |
| *Lithosiini gensp CRS7* | Erebidae | incidental |  | 58480324 |
| *Lithosiini gensp. Blackgold* | Erebidae | 0.52 |  | 33123758 |
| *Lithosiini gensp. CRS1* | Erebidae | 0.52 |  | 54221849 |
| *Lithosiini gensp. CRS2* | Erebidae | 0.52 |  | 56289600 |
| *Lithosiini gensp. CRS3* | Erebidae | 1.57 |  | 56370375 |
| *Lithosiini gensp. CRS4* | Erebidae | 0.52 |  | 56728650 |
| *Lithosiini gensp. CRS5* | Erebidae | 0.52 |  | 49808273 |
| *Lithosiini gensp. CRS6* | Erebidae | 1.57 |  | 56618476 |
| *Luceria oculalis* | Erebidae | 6.28 |  | 31844790 |
| *Luceria sp. CRS1* | Erebidae | 11.52 |  | 56928173 |
| *Lysimelia lenis* | Erebidae | incidental |  | 99702880 |
| *Macaduma aff. toxophora* | Erebidae | incidental |  | 59108463 |
| *Macaduma strongyla* | Erebidae | incidental |  | 59534224 |
| *Metaphoenia rhodias* | Erebidae | 0.52 |  | 56618453 |
| *Meyrickella ruptellus* | Erebidae | 0.52 |  | 56923816 |
| *Meyrickella torquesauria* | Erebidae | 1.05 |  | 50773384 |
| *Microstola ammoscia* | Erebidae | 3.66 |  | 56618470 |
| *Miltochrista pyraula* | Erebidae | 2.62 |  | 58112241 |
| *Mocis alterna* | Erebidae | 1.05 |  | 50338665 |
| *Mocis frugalis* | Erebidae | 18.32 |  | 30918317 |
| *Mocis trifasciata* | Erebidae | 8.9 |  | 32401121 |
| *Naarda xanthonephra* | Erebidae | 5.24 |  | 55466678 |
| *Nodaria aneliopis* | Erebidae | 1.57 |  | 31780441 |
| *Nodaria cornicalis* | Erebidae | 3.66 |  | 31152534 |
| *Notata modicus* | Erebidae | 15.18 |  | 31465080 |
| *Nudaria mollis* | Erebidae | 14.66 |  | 54707505 |
| *Nyctemera baulus* | Erebidae | 13.09 |  | 30501902 |
| *Oeonistis altica* | Erebidae | 0.52 |  | 51211042 |
| *Olene mendosa* | Erebidae | 0.52 |  | 54907872 |
| *Ophiusa disjungens* | Erebidae | 0.52 |  | 56327480 |
| *Ophyx eurrhoa* | Erebidae | 0.52 |  | 52345690 |
| *Orgyia sp. CRS1* | Erebidae | 0.52 |  | 51669385 |
| *Oruza cariosa* | Erebidae | 1.05 |  | 49407536 |
| *Oxyodes sp. CRS1* | Erebidae | 0.52 |  | 56466103 |
| *Oxyodes tricolor* | Erebidae | 0.52 |  | 48898478 |
| *Pantydia capistrata* | Erebidae | 0.52 |  | 31203911 |
| *Pantydia metaspila* | Erebidae | 0.52 |  | 51211059 |
| *Pantydia sparsa* | Erebidae | 0.52 |  | 50621777 |
| *Pherechoa crypsichlora* | Erebidae | 5.24 |  | 31780424 |
| *Pherechoa sp. CRS1* | Erebidae | 1.57 |  | 55362892 |
| *Philenora aspectalella* | Erebidae | 7.85 |  | 56924934 |
| *Philenora chionastis* | Erebidae | 3.66 |  | 33125087 |
| *Philogethes metableta* | Erebidae | 0.52 |  | 56327481 |
| *Phyllodes imperialis* | Erebidae | 0.52 |  | 52572769 |
| *Phytometra laevis* | Erebidae | 4.19 |  | 31844796 |
| *Praxis marmarinopa* | Erebidae | 4.19 |  | 30806615 |
| *Progonia sp. EF02 in BOLD* | Erebidae | 8.9 |  | 54707545 |
| *Progonia umbrifera* | Erebidae | 11.52 |  | 55224278 |
| *Rivula biagi* | Erebidae | 1.05 |  | 56289593 |
| *Rivula curvifera* | Erebidae | 0.52 |  | 31605665 |
| *Rusicada revocans* | Erebidae | 0.52 |  | 54112594 |
| *Sandava xylistis* | Erebidae | 1.05 |  | 33125076 |
| *Saroba trimaculata* | Erebidae | 0.52 |  | 40814124 |
| *Scaptesyle dichotoma* | Erebidae | 1.57 |  | 56924937 |
| *Schistophleps albida* | Erebidae | 5.76 |  | 50671011 |
| *Schistorhynx unistriga* | Erebidae | 0.52 |  | 53792881 |
| *Serrodes campana* | Erebidae | 1.57 |  | 50338666 |
| *Simplicia sp. CRS1* | Erebidae | 0.52 |  | 54510050 |
| *Sommeria marmorea* | Erebidae | 0.52 |  | 56370367 |
| *Speiredonia spectans* | Erebidae | 0.52 |  | 39961965 |
| *Spilosoma curvata* | Erebidae | 3.14 |  | 31465078 |
| *Symmetrodes sciocosma* | Erebidae | 32.98 |  | 30501903 |
| *Sympis parkeri* | Erebidae | 0.52 |  | 50247996 |
| *Sympis rufibasis* | Erebidae | 0.52 |  | 54907881 |
| *Teulisna bipunctata* | Erebidae | incidental |  | 59857580 |
| *Thallarcha epileuca* | Erebidae | incidental |  | 60024754 |
| *Thallarcha levis* | Erebidae | 0.52 |  | 53593764 |
| *Thallarcha sp. CRS1* | Erebidae | 0.52 |  | 56023529 |
| *Tolpia conscitulana* | Erebidae | 5.76 |  | 49097027 |
| *Trigonistis demonias* | Erebidae | 1.05 |  | 50621733 |
| *Trigonodes hyppasia* | Erebidae | 6.81 |  | 30918306 |
| *Utetheisa aegrotum* | Erebidae | 1.05 |  | 45858866 |
| *Utetheisa pulchelloides* | Erebidae | 5.76 |  | 32883931 |
| *Euteliidae gensp. CRS1* | Euteliidae | 0.52 |  | 55862436 |
| *Lophoptera aff. melanisigera-vittigera* | Euteliidae | 0.52 |  | 48898561 |
| *Lophoptera hemithyris* | Euteliidae | 1.57 |  | 49307817 |
| *Lophoptera nama* | Euteliidae | 0.52 |  | 56924950 |
| *Lophoptera sp. CRS1* | Euteliidae | 2.09 |  | 53433574 |
| *Paectes cyanodes* | Euteliidae | incidental |  | 57095049 |
| *Pataeta carbo* | Euteliidae | 1.05 |  | 50038643 |
| *Penicillaria jocosatrix* | Euteliidae | 1.05 |  | 55859448 |
| *Stictoptera cucullioides* | Euteliidae | incidental |  | 57095060 |
| *Targalla delatrix* | Euteliidae | 2.62 |  | 56272193 |
| *Targalla scelerata* | Euteliidae | 0.52 |  | 50343366 |
| *Anarsia anisodonta* | Gelechiidae | 0.52 |  | 51569163 |
| *Aproaerema isoscelixantha* | Gelechiidae | 1.05 |  | 56312827 |
| *Ardozyga sp CRS1* | Gelechiidae | 0.52 |  | 51470214 |
| *Dichomeridinae gensp. CRS1* | Gelechiidae | 2.09 |  | 48898500 |
| *Dichomeridinae gensp. CRS2* | Gelechiidae | 0.52 |  | 57013856 |
| *Dichomeris acuminatus* | Gelechiidae | 2.09 |  | 48898483 |
| *Dichomeris sp. CRS1* | Gelechiidae | 0.52 |  | 56577862 |
| *Dichomeris sp. CRS2* | Gelechiidae | 0.52 |  | 54707536 |
| *Gelechiidae aff. Thiotricha* | Gelechiidae | 1.57 |  | 53792894 |
| *Gelechiidae gensp. CRS1* | Gelechiidae | 0.52 |  | 54238570 |
| *Gelechiidae gensp. CRS2* | Gelechiidae | 0.52 |  | 52915134 |
| *Gelechiidae gensp. CRS3* | Gelechiidae | 0.52 |  | 51554201 |
| *Hypatima aff. deviella* | Gelechiidae | 0.52 |  | 50773386 |
| *Hypatima cyrtopleura* | Gelechiidae | 1.05 |  | 55861686 |
| *Hypatima discissa* | Gelechiidae | 0.52 |  | 49808280 |
| *Hypatima sp. CRS1* | Gelechiidae | 0.52 |  | 56370338 |
| *Idiophantis habrias* | Gelechiidae | 0.52 |  | 55989617 |
| *Stegasta variana* | Gelechiidae | 0.52 |  | 55861700 |
| *Thiotricha atractodes* | Gelechiidae | 24.61 |  | 46911886 |
| *Thiotricha gensp. CRS1* | Gelechiidae | incidental |  | 59928441 |
| *Thiotricha margarodes* | Gelechiidae | incidental |  | 59419178 |
| *Abraxas expectata* | Geometridae | 1.05 |  | 46911885 |
| *Aeolochroma turneri* | Geometridae | 6.28 |  | 50621742 |
| *Aeolochroma viridicata* | Geometridae | 0.52 |  | 48847961 |
| *Agathia pisina* | Geometridae | 3.14 |  | 54208246 |
| *Agathia prasinaspis* | Geometridae | 0.52 |  | 39961825 |
| *Antimimistis attenuata* | Geometridae | 5.24 |  | 48898494 |
| *Boarmiini gensp. CRS1* | Geometridae | 0.52 |  | 53207372 |
| *Boarmiini gensp. CRS2* | Geometridae | 0.52 |  | 48898416 |
| *Borbacha euchrysa* | Geometridae | incidental |  | 61894716 |
| *Bracca rotundata* | Geometridae | 5.76 |  | 31340574 |
| *Calluga costalis* | Geometridae | 6.81 |  | 33123760 |
| *Capusa sp. CRS1* | Geometridae | 4.71 |  | 49808286 |
| *Casbia albinotata* | Geometridae | 0.52 |  | 50276524 |
| *Casbia calliorma* | Geometridae | 4.19 |  | 50765334 |
| *Casbia rectaria* | Geometridae | 11.52 |  | 50023775 |
| *Casbia scardamiata* | Geometridae | 3.14 |  | 50271559 |
| *Casbia sp. CRS1* | Geometridae | 0.52 |  | 45858829 |
| *Casbia sp. CRS2* | Geometridae | 2.62 |  | 33125037 |
| *Casbia sp. CRS3* | Geometridae | 0.52 |  | 49307835 |
| *Casbia sp. CRS4* | Geometridae | 0.52 |  | 53009869 |
| *Catoria delectaria* | Geometridae | 0.52 |  | 50031216 |
| *Chaetolopha emporias* | Geometridae | incidental |  | 60184089 |
| *Chiasmia tessellata* | Geometridae | 0.52 |  | 50131500 |
| *Chloroclystis approximata* | Geometridae | 1.05 |  | 56728658 |
| *Chloroclystis elaeopa* | Geometridae | 2.62 |  | 53016862 |
| *Chloroclystis mniochroa* | Geometridae | 1.05 |  | 55466661 |
| *Chloroclystis perissa* | Geometridae | 2.09 |  | 50765322 |
| *Chorodna strixaria* | Geometridae | 3.14 |  | 49924272 |
| *Chrysocraspeda sp. ANIC5* | Geometridae | 1.05 |  | 32633404 |
| *Circopetes obtusata* | Geometridae | 0.52 |  | 38377456 |
| *Cleora illustraria* | Geometridae | 0.52 |  | 50886061 |
| *Cleora repetita* | Geometridae | 2.09 |  | 48898451 |
| *Collix ghosha* | Geometridae | 2.62 |  | 43305123 |
| *Comibaena inductaria* | Geometridae | 1.05 |  | 54341093 |
| *Comibaena mariae* | Geometridae | 1.05 |  | 49407526 |
| *Comostola aff. subtiliaria* | Geometridae | 1.05 |  | 41027894 |
| *Comostola citrolimbaria* | Geometridae | incidental |  | 41251705 |
| *Comostola laesaria* | Geometridae | 0.52 |  | 42306734 |
| *Comostola leucomerata* | Geometridae | 1.05 |  | 30918311 |
| *Comostola pyrrhogona* | Geometridae | 1.57 |  | 56289582 |
| *Cosmogonia decorata* | Geometridae | 6.81 |  | 50199295 |
| *Crasilogia gressitti* | Geometridae | 2.62 |  | 31152528 |
| *Craspedosis leucosticta* | Geometridae | incidental |  | 40547086 |
| *Cyclophora frenaria* | Geometridae | 1.05 |  | 50028727 |
| *Cyclophora obstataria* | Geometridae | 1.05 |  | 55873544 |
| *Dysphania numana* | Geometridae | 0.52 |  | 30857345 |
| *Ectropis aff. gravis* | Geometridae | 0.52 |  | 56272203 |
| *Ectropis bhurmitra* | Geometridae | 0.52 |  | 54694868 |
| *Ectropis bispinaria* | Geometridae | 2.09 |  | 50735976 |
| *Ennominae gensp. CRS1* | Geometridae | 0.52 |  | 56466067 |
| *Ennominae gensp. CRS2* | Geometridae | 0.52 |  | 32264626 |
| *Ennominae gensp. CRS3* | Geometridae | 0.52 |  | 49412451 |
| *Eois cymatodes* | Geometridae | 3.14 |  | 32878620 |
| *Epidesmia phoenicina* | Geometridae | 0.52 |  | 56618495 |
| *Epidesmia reservata* | Geometridae | incidental |  | 58318049 |
| *Epyaxa sodaliata* | Geometridae | incidental |  | 59948189 |
| *Eucyclodes fascinans* | Geometridae | 2.09 |  | 43305087 |
| *Eumelea rosalia* | Geometridae | 0.52 |  | 50338639 |
| *Geometridae Emerald* | Geometridae | 0.52 |  | 49825643 |
| *Geometridae gensp. aff Nearcha* | Geometridae | 0.52 |  | 55976831 |
| *Geometridae gensp. CRS1* | Geometridae | 0.52 |  | 56466074 |
| *Geometridae gensp. CRS2* | Geometridae | 0.52 |  | 30918308 |
| *Geometridae gensp. CRS3* | Geometridae | 0.52 |  | 52915146 |
| *Geometridae gensp. CRS4* | Geometridae | 0.52 |  | 50247998 |
| *Geometridae gensp. CRS5* | Geometridae | 0.52 |  | 52204505 |
| *Geometridae gensp. CRS6* | Geometridae | 0.52 |  | 49808176 |
| *Geometridae gensp. CRS7* | Geometridae | incidental |  | 58318047 |
| *Geometridae gensp. CRS8* | Geometridae | incidental |  | 59217526 |
| *Geometridae WhiteWingsBrownCosta* | Geometridae | 3.14 |  | 56272204 |
| *Gymnoscelis callichlora* | Geometridae | 0.52 |  | 54707558 |
| *Gymnoscelis ischnophylla* | Geometridae | 1.05 |  | 48974177 |
| *Gymnoscelis lophopus* | Geometridae | 3.14 |  | 31844765 |
| *Gymnoscelis mesophoena* | Geometridae | 0.52 |  | 31844789 |
| *Hypodoxa emiliaria* | Geometridae | 0.52 |  | 31605650 |
| *Hyposidra incomptaria* | Geometridae | 3.14 |  | 55362883 |
| *Idaea argophylla* | Geometridae | incidental |  | 60184087 |
| *Idaea coercita* | Geometridae | 2.62 |  | 49523555 |
| *Idaea sp. CRS1* | Geometridae | 0.52 |  | 51470201 |
| *Idiodes apicata* | Geometridae | 1.57 |  | 33125085 |
| *Idiodes sp. CRS1* | Geometridae | 0.52 |  | 49825641 |
| *Idioides aff. ANIC2* | Geometridae | 1.05 |  | 31605655 |
| *Larentiinae aff Visiana* | Geometridae | 2.62 |  | 55861692 |
| *Lophosigna catasticta* | Geometridae | incidental |  | 62842287 |
| *Luxiaria ochrophara* | Geometridae | 4.19 |  | 46358674 |
| *Maxates orthodesma* | Geometridae | 2.09 |  | 31605637 |
| *Metallochlora militaris* | Geometridae | 1.57 |  | 51211040 |
| *Metallochlora venusta* | Geometridae | incidental |  | 59314630 |
| *Mnesiloba eupitheciata* | Geometridae | 1.57 |  | 31203905 |
| *Nadagara xylotrema* | Geometridae | 1.05 |  | 56924947 |
| *Nearcha sp. CRS1* | Geometridae | 0.52 |  | 54694832 |
| *Neogyne elongata* | Geometridae | incidental |  | 61568459 |
| *Oenochlora imperialis* | Geometridae | incidental |  | 97522907 |
| *Oenochroma lissoscia* | Geometridae | incidental |  | 58115647 |
| *Oenochroma turneri* | Geometridae | 0.52 |  | 49412233 |
| *Oenochroma vetustaria* | Geometridae | 0.52 |  | 50243696 |
| *Organopoda olivescens* | Geometridae | 0.52 |  | 50023774 |
| *Paradromulia ambigua* | Geometridae | 3.14 |  | 53792874 |
| *Parepisparis pallidus* | Geometridae | incidental |  | 97597728 |
| *Perixera flavirubra* | Geometridae | 1.05 |  | 50550156 |
| *Perixera obliviaria* | Geometridae | 1.05 |  | 47382235 |
| *Perixera punctata* | Geometridae | 1.05 |  | 50199299 |
| *Phrissogonus laticostata* | Geometridae | 0.52 |  | 31844783 |
| *Pingasa chlora* | Geometridae | 0.52 |  | 33123756 |
| *Poecilasthena pulchraria* | Geometridae | 0.52 |  | 32332146 |
| *Poecilasthena Rusty* | Geometridae | 1.05 |  | 56289566 |
| *Polyacme dissimilis* | Geometridae | 3.66 |  | 53433584 |
| *Polyclysta hypogrammata* | Geometridae | 0.52 |  | 56583687 |
| *Prasinocyma albicosta* | Geometridae | 1.57 |  | 50669057 |
| *Prasinocyma caniola* | Geometridae | 0.52 |  | 49421958 |
| *Prasinocyma floresaria* | Geometridae | 2.09 |  | 31844782 |
| *Prasinocyma lychnopasta* | Geometridae | 1.05 |  | 53792879 |
| *Probithia sp. ANIC1* | Geometridae | 0.52 |  | 50243691 |
| *Probithia sp. CRS1* | Geometridae | 0.52 |  | 56023513 |
| *Protuliocnemis biplagiata* | Geometridae | 0.52 |  | 43305164 |
| *Protuliocnemis partita* | Geometridae | 0.52 |  | 55695057 |
| *Racotis maculata* | Geometridae | 0.52 |  | 50108637 |
| *Sauris plumipes* | Geometridae | 1.05 |  | 54495240 |
| *Scardamia ithyzona* | Geometridae | 1.05 |  | 55980661 |
| *Scopula sp. CRS1* | Geometridae | 4.71 |  | 50621757 |
| *Scopula sp. CRS2* | Geometridae | 1.57 |  | 31203899 |
| *Scopula sublinearia* | Geometridae | 4.19 |  | 54694826 |
| *Selidosema agoraea* | Geometridae | 1.57 |  | 49907477 |
| *Sigilliclystis insigillata* | Geometridae | 2.09 |  | 49421961 |
| *Sterrhinae gensp. CRS1* | Geometridae | 0.52 |  | 52927295 |
| *Sterrhinae gensp. CRS2* | Geometridae | incidental |  | 57095061 |
| *Sterrhinae gensp. CRS3* | Geometridae | incidental |  | 58563323 |
| *Sterrhinae gensp. CRS4* | Geometridae | incidental |  | 57095059 |
| *Thalassodes pilaria* | Geometridae | 2.09 |  | 50108631 |
| *Traminda aventiaria* | Geometridae | 5.76 |  | 33123763 |
| *Glyphipterix marmaropa* | Glyphipterigidae | 4.19 | Yes | 50202209 |
| *Acrocercops aff. hedymopa* | Gracillariidae | Incidental |  | 57102507 |
| *Acrocercops aff. macaria* | Gracillariidae | 1.05 |  | 56466061 |
| *Acrocercops chionosema* | Gracillariidae | 0.52 |  | 53555323 |
| *Acrocercops hedymopa* | Gracillariidae | 0.52 |  | 55362872 |
| *Acrocercops hoplocala* | Gracillariidae | 0.52 |  | 54694812 |
| *Acrocercops macaria* | Gracillariidae | 0.52 |  | 33125084 |
| *Acrocercops plebeia* | Gracillariidae | 2.09 |  | 54707538 |
| *Acrocercops sp. CRS1* | Gracillariidae | 0.52 |  | 51441231 |
| *Acrocercops sp. CRS2* | Gracillariidae | 0.52 |  | 54886212 |
| *Caloptilia euglypta* | Gracillariidae | 3.14 |  | 54907783 |
| *Caloptilia sp. ANIC 1* | Gracillariidae | 0.52 |  | 56289575 |
| *Caloptilia sp. CRS1* | Gracillariidae | incidental |  | 100135966 |
| *Caloptilia xanthopharella* | Gracillariidae | incidental |  | 59884494 |
| *Epicephala albistriatella* | Gracillariidae | 0.52 |  | 56928168 |
| *Epicephala sp. CRS1* | Gracillariidae | 0.52 |  | 55224283 |
| *Epicephala sp. CRS2* | Gracillariidae | incidental |  | 60190448 |
| *Gibbovalva quadrifasciata* | Gracillariidae | 0.52 |  | 54022464 |
| *Gracillariidae gensp. CRS6* | Gracillariidae | incidental |  | 59885655 |
| *Gracillariidae sp. 1* | Gracillariidae | 0.52 |  | 55224199 |
| *Gracillariidae sp. 2* | Gracillariidae | 0.52 |  | 54495201 |
| *Gracillariidae sp. 3* | Gracillariidae | 1.05 |  | 56004698 |
| *Gracillariidae sp. 4* | Gracillariidae | 0.52 |  | 54694809 |
| *Gracillariidae sp. 5* | Gracillariidae | 0.52 |  | 53207519 |
| *Neurostrota gunniella* | Gracillariidae | 0.52 |  | 56618464 |
| *Phyllocnistis citrella* | Gracillariidae | 0.52 |  | 53207339 |
| *Phyllocnistis sp. CRS1* | Gracillariidae | 0.52 |  | 55224217 |
| *Phyllocnistis sp. CRS2* | Gracillariidae | 1.05 |  | 51554197 |
| *Phyllocnistis sp. CRS3* | Gracillariidae | 0.52 |  | 56016981 |
| *Heliozela sp. CRS1* | Heliozelidae | incidental |  | 59524496 |
| *Oncopera brachyphylla* | Hepialidae | 0.52 |  | 48898431 |
| *Oncopera mitocera* | Hepialidae | 1.05 |  | 50023952 |
| *Oxycanus buluwandji* | Hepialidae | 4.71 |  | 31096104 |
| *Lactura calliphylla* | Lacturidae | incidental |  | 61568458 |
| *Lactura panopsia* | Lacturidae | 0.52 |  | 50028734 |
| *Lasiocampidae gensp. CRS1* | Lasiocampidae | 0.52 |  | 56583661 |
| *Porela aff. dilineata* | Lasiocampidae | 0.52 |  | 54907803 |
| *Crocanthes halurga* | Lecithoceridae | 1.05 |  | 56583680 |
| *Crocanthes prasinopis* | Lecithoceridae | 1.57 |  | 53009859 |
| *Crocanthes trizona* | Lecithoceridae | incidental |  | 59217531 |
| *Lecithocera cyamitis* | Lecithoceridae | 2.09 |  | 56728632 |
| *Lecithocera imprudens* | Lecithoceridae | 3.14 |  | 52915151 |
| *Lecithoceridae gensp. CRS1* | Lecithoceridae | 1.05 |  | 49917914 |
| *Lecithoceridae gensp. CRS2* | Lecithoceridae | 0.52 |  | 53207566 |
| *Lecithoceridae gensp. CRS3* | Lecithoceridae | incidental |  | 59885656 |
| *Anaxidia lozogramma* | Limacodidae | incidental |  | 58100716 |
| *Doratifera pinguis* | Limacodidae | 1.05 |  | 33125056 |
| *Limacodidae gensp. CRS1* | Limacodidae | 0.52 |  | 45858835 |
| *Praesusica placerodes* | Limacodidae | 0.52 |  | 55980660 |
| *Squamosa barymorpha* | Limacodidae | 0.52 |  | 48847966 |
| *Euproctis aganopa* | Lymantriidae | incidental |  | 59109103 |
| *Lymantriinae gensp CRS1* | Lymantriidae | 2.09 |  | 33125077 |
| *Lyonetiidae gensp. CRS1* | Lyonetiidae | incidental |  | 60188639 |
| *Stegommata leptomitella* | Lyonetiidae | incidental |  | 60081671 |
| *Tasmantrix thula* | Micropterigidae | 12.57 |  | 51470227 |
| *Nepticulidae gensp. CRS1* | Nepticulidae | 0.52 |  | 56016955 |
| *Nepticulidae gensp. CRS2* | Nepticulidae | 0.52 |  | 53207514 |
| *Acronicta psorallina* | Noctuidae | incidental |  | 58115642 |
| *Adisura marginalis* | Noctuidae | 1.57 |  | 49412234 |
| *Aedia olivescens* | Noctuidae | 0.52 |  | 46911857 |
| *Agrotis munda* | Noctuidae | 1.57 |  | 32401160 |
| *Amyna axis* | Noctuidae | 1.05 |  | 48974126 |
| *Amyna natalis* | Noctuidae | 1.05 |  | 54572371 |
| *Athetis maculatra* | Noctuidae | 6.81 |  | 53335580 |
| *Athetis sp. CRS1* | Noctuidae | 0.52 |  | 55873536 |
| *Australothis rubrescens* | Noctuidae | 2.62 |  | 31605669 |
| *Callopistria sp. CRS1* | Noctuidae | 0.52 |  | 48898498 |
| *Chrysodeixis acuta* | Noctuidae | 1.57 |  | 50038642 |
| *Chrysodeixis eriosoma* | Noctuidae | 0.52 |  | 50338657 |
| *Chrysodeixis illuminata* | Noctuidae | 1.57 |  | 50131505 |
| *Chrysodeixis sp. CRS1* | Noctuidae | 0.52 |  | 32878512 |
| *Chrysodeixis sp. CRS2* | Noctuidae | 0.52 |  | 50024921 |
| *Chrysodeixis sp. CRS3* | Noctuidae | 0.52 |  | 43299664 |
| *Chrysodeixis sp. CRS4* | Noctuidae | 0.52 |  | 52999340 |
| *Condica dolorosa* | Noctuidae | 0.52 |  | 51669374 |
| *Condica illecta* | Noctuidae | 2.09 |  | 47382229 |
| *Cosmodes elegans* | Noctuidae | 2.09 |  | 30806626 |
| *Ctenoplusia albostriata* | Noctuidae | 1.05 |  | 50105029 |
| *Cycloprora nodyna* | Noctuidae | 1.05 |  | 50004068 |
| *Elusa oenolopha* | Noctuidae | 1.05 |  | 56583713 |
| *Elusa semipecten* | Noctuidae | 0.52 |  | 54907853 |
| *Elusa sp. CRS1* | Noctuidae | 1.05 |  | 56618435 |
| *Helicoverpa armigera* | Noctuidae | 1.05 |  | 56583688 |
| *Helicoverpa assulta* | Noctuidae | 1.05 |  | 50131502 |
| *Helicoverpa punctigera* | Noctuidae | 0.52 |  | 54499936 |
| *Heliothinae gensp. CRS1* | Noctuidae | 0.52 |  | 31203908 |
| *Heliothinae gensp. CRS2* | Noctuidae | 0.52 |  | 50023792 |
| *Heliothis punctifera* | Noctuidae | 0.52 |  | 50145234 |
| *Hypobleta cymaea* | Noctuidae | 2.09 |  | 56289564 |
| *Leucania abdominalis* | Noctuidae | 3.66 |  | 31281844 |
| *Leucania dasycnema* | Noctuidae | 1.05 |  | 30806630 |
| *Leucania designata* | Noctuidae | 3.14 |  | 49532766 |
| *Leucania leucosta* | Noctuidae | 6.81 |  | 52136111 |
| *Leucania linearis* | Noctuidae | 0.52 |  | 55377986 |
| *Leucania loreyi* | Noctuidae | 1.05 |  | 56312815 |
| *Leucania polysticha* | Noctuidae | 0.52 |  | 30806617 |
| *Leucania porphyrodes* | Noctuidae | 0.52 |  | 56466053 |
| *Leucania stenographa* | Noctuidae | 1.57 |  | 32401100 |
| *Leucania yu* | Noctuidae | 2.09 |  | 49663012 |
| *Leucogonia ekeikei* | Noctuidae | incidental |  | 60024751 |
| *Maliattha ritsemae* | Noctuidae | 0.52 |  | 50199289 |
| *Maliattha signifera* | Noctuidae | 1.57 |  | 47382239 |
| *Mythimna formosana* | Noctuidae | 1.57 |  | 53207537 |
| *Mythimna sp. CRS1* | Noctuidae | 0.52 |  | 45858840 |
| *Noctuidae aff Athetis* | Noctuidae | 0.52 |  | 55373469 |
| *Noctuidae gensp. CRS1* | Noctuidae | 0.52 |  | 50031210 |
| *Noctuidae gensp. CRS2* | Noctuidae | 0.52 |  | 49917912 |
| *Noctuidae gensp. CRS3* | Noctuidae | 0.52 |  | 32878612 |
| *Noctuidae gensp. CRS4* | Noctuidae | 0.52 |  | 43305113 |
| *Noctuidae gensp. CRS5* | Noctuidae | 0.52 |  | 50338654 |
| *Noctuidae gensp. CRS6* | Noctuidae | 0.52 |  | 52434109 |
| *Pachythrix hampsoni* | Noctuidae | incidental |  | 60059606 |
| *Parapadna zonophora* | Noctuidae | 0.52 |  | 55861685 |
| *Plusiinae gensp. CRS1* | Noctuidae | 0.52 |  | 33123757 |
| *Plusiinae gensp. CRS2* | Noctuidae | 2.09 |  | 33123749 |
| *Proteuxoa hypochalchis* | Noctuidae | 1.05 |  | 53207397 |
| *Proteuxoa tibiata* | Noctuidae | 0.52 |  | 51470193 |
| *Spodoptera litura* | Noctuidae | 5.24 |  | 31464708 |
| *Spodoptera mauritia* | Noctuidae | 4.19 |  | 56928169 |
| *Spodoptera sp. CRS1* | Noctuidae | 0.52 |  | 56299161 |
| *Squamipalpis pantoea* | Noctuidae | 0.52 |  | 55224229 |
| *Thysanoplusia lectula* | Noctuidae | 1.05 |  | 48898428 |
| *Thysanoplusia orichalcea* | Noctuidae | 1.05 |  | 31464731 |
| *Xanthodes congenita* | Noctuidae | 0.52 |  | 50338661 |
| *Xanthodes transversa* | Noctuidae | 0.52 |  | 41359309 |
| *Acatapaustus atrinota* | Nolidae | 2.62 |  | 49085731 |
| *Acatapaustus leucospila* | Nolidae | 1.57 |  | 43299706 |
| *Acatapaustus mesoleuca* | Nolidae | 6.28 |  | 50765340 |
| *Cacyparis brevipennis* | Nolidae | incidental |  | 69409881 |
| *Cacyparis melanolitha* | Nolidae | incidental |  | 60184086 |
| *Calathusa hypotherma* | Nolidae | 0.52 |  | 54886227 |
| *Calathusa sp CRS unsorted* | Nolidae | 2.09 |  | 56928162 |
| *Earias chlorodes* | Nolidae | 1.05 |  | 52915147 |
| *Earias flavida* | Nolidae | 1.57 |  | 30918324 |
| *Earias luteolaria* | Nolidae | 0.52 |  | 51211033 |
| *Earias smaragdina* | Nolidae | 2.09 |  | 40814348 |
| *Earias vittella* | Nolidae | 1.57 |  | 50023780 |
| *Etanna basalis* | Nolidae | 1.05 |  | 54907857 |
| *Etanna breviuscula* | Nolidae | 3.66 |  | 56683806 |
| *Gadirtha inexacta* | Nolidae | 1.05 |  | 50874571 |
| *Garella curiosa* | Nolidae | 0.52 |  | 56583736 |
| *Maceda mansueta* | Nolidae | 18.85 |  | 56272219 |
| *Maceda rotundimacula* | Nolidae | 0.52 |  | 54907862 |
| *Maurilia iconica* | Nolidae | incidental |  | 62842253 |
| *Negeta contrariata* | Nolidae | 0.52 |  | 53035502 |
| *Nola bifascialis* | Nolidae | 2.62 |  | 31203917 |
| *Nola desmotes* | Nolidae | 4.19 |  | 53247982 |
| *Nola elaphra* | Nolidae | 0.52 |  | 49287982 |
| *Nola epicentra* | Nolidae | 0.52 |  | 52623775 |
| *Nola fasciata* | Nolidae | 3.66 |  | 51470213 |
| *Nola fraterna* | Nolidae | 1.05 |  | 50621753 |
| *Nola pygmaeodes* | Nolidae | 8.38 |  | 55224208 |
| *Nola sp. CRS1* | Nolidae | 0.52 |  | 31844795 |
| *Nola sp. CRS2* | Nolidae | 0.52 |  | 54707546 |
| *Nola sp. CRS3* | Nolidae | 0.52 |  | 52879162 |
| *Nola sp. CRS4* | Nolidae | 0.52 |  | 55465213 |
| *Nola sphaerospila* | Nolidae | 4.19 |  | 54886178 |
| *Nola taeniata* | Nolidae | 3.66 |  | 55483233 |
| *Nola tornotis* | Nolidae | 1.05 |  | 49307833 |
| *Nolidae gensp. CRS1* | Nolidae | incidental |  | 58480326 |
| *Nycteola indica* | Nolidae | 1.05 |  | 56016979 |
| *Nycteola polycyma* | Nolidae | 1.05 |  | 33125079 |
| *Nycteola symmicta* | Nolidae | 0.52 |  | 33125029 |
| *Ochthophora sericina* | Nolidae | incidental |  | 61786611 |
| *Paracrama latimargo* | Nolidae | incidental |  | 68856077 |
| *Pisara hyalospila* | Nolidae | 23.56 |  | 48898452 |
| *Risoba obstructa* | Nolidae | 0.52 |  | 39399663 |
| *Sarrothripinae gensp. CRS1* | Nolidae | 0.52 |  | 56004671 |
| *Sarrothripinae gensp. CRS2* | Nolidae | 1.05 |  | 56299165 |
| *Thriponea orbiculigera* | Nolidae | 0.52 |  | 52346192 |
| *Uraba lugens* | Nolidae | 6.28 |  | 55692142 |
| *Westermannia gloriosa* | Nolidae | 0.52 |  | 50343371 |
| *Cascera bella* | Notodontidae | 1.05 |  | 50671002 |
| *Destolmia lineata* | Notodontidae | 2.09 |  | 32264404 |
| *Gargettiana punctatissima* | Notodontidae | 1.05 |  | 49808208 |
| *Heterocampinae gensp. CRS1* | Notodontidae | 0.52 |  | 50105022 |
| *Kamalia multipunctata* | Notodontidae | 0.52 |  | 50669037 |
| *Lasioceros aroa* | Notodontidae | 1.05 |  | 54228799 |
| *Neostauropus viridissimus* | Notodontidae | 1.57 |  | 49723565 |
| *Notodontidae gensp. CRS1* | Notodontidae | 0.52 |  | 50669047 |
| *Notodontidae gensp. CRS2* | Notodontidae | 0.52 |  | 51211060 |
| *Notodontidae gensp. CRS3* | Notodontidae | 0.52 |  | 53792884 |
| *Ochrogaster lunifer* | Notodontidae | 3.14 |  | 32462102 |
| *Omichlis hadromeres* | Notodontidae | 2.09 |  | 31780435 |
| *Ortholomia moluccana* | Notodontidae | 0.52 |  | 50036611 |
| *Pheraspis mesotypa* | Notodontidae | 0.52 |  | 50124327 |
| *Olbonoma sp. CRS1* | Oechophoridae | incidental |  | 61786590 |
| *Garrha costimacula* | Oecophidae | incidental |  | 59217534 |
| *Garrha sp. CRS1* | Oecophidae | incidental |  | 61569592 |
| *Acantholena siccella* | Oecophoridae | 0.52 |  | 55900090 |
| *Acorotricha crystanta* | Oecophoridae | 0.52 |  | 33125074 |
| *Barea sp. CRS1* | Oecophoridae | incidental |  | 55861703 |
| *Callithauma pyrites* | Oecophoridae | 0.52 | Yes | 50621729 |
| *Chezala privatella* | Oecophoridae | 3.14 |  | 31464728 |
| *Elaphromorpha axierasta* | Oecophoridae | incidental | Yes | 61786600 |
| *Elaphromorpha glycymilicha* | Oecophoridae | incidental | Yes | 60190440 |
| *Erythrisa oenoessa* | Oecophoridae | 0.52 | Yes | 54243543 |
| *Eulechria aff. malacostola* | Oecophoridae | incidental |  | 59955701 |
| *Eulechria sp. CRS1* | Oecophoridae | 0.52 |  | 55124318 |
| *Locheutis auchmera* | Oecophoridae | incidental |  | 59524487 |
| *Oecophoridae gensp. CRS1* | Oecophoridae | 0.52 |  | 55861703 |
| *Oecophoridae gensp. CRS2* | Oecophoridae | 1.05 |  | 54495219 |
| *Oecophoridae gensp. CRS3* | Oecophoridae | 1.05 |  | 54341120 |
| *Oecophoridae gensp. CRS4* | Oecophoridae | 0.52 |  | 53555334 |
| *Oecophoridae gensp. CRS5* | Oecophoridae | 0.52 |  | 56923831 |
| *Oecophoridae gensp. CRS6* | Oecophoridae | incidental |  | 58480352 |
| *Oecophorinae sp. CRS1* | Oecophoridae | 0.52 |  | 50202202 |
| *Philobota aff mathematica* | Oecophoridae | 6.81 |  | 54238551 |
| *Philobota baryptera* | Oecophoridae | 3.66 |  | 56618496 |
| *Philobota sp. CRS1* | Oecophoridae | 0.52 |  | 33125023 |
| *Philobota sp. CRS2* | Oecophoridae | incidental |  | 58480344 |
| *Philobota sp. CRS3* | Oecophoridae | 2.09 |  | 56916523 |
| *Philobota sp. CRS4* | Oecophoridae | 3.66 |  | 51856690 |
| *Psaroxantha aff. ANIC2* | Oecophoridae | 0.52 |  | 58399276 |
| *Tachystola hemisema* | Oecophoridae | 0.52 |  | 52434120 |
| *Tachystola thiasotis* | Oecophoridae | 0.52 | Yes | 56327475 |
| *Tortricopsis pyroptis* | Oecophoridae | incidental |  | 59109102 |
| *Opostega aff. nubifera* | Opostegidae | 0.52 |  | 31780459 |
| *Opostega GoldMink* | Opostegidae | 0.52 |  | 56466100 |
| *Opostega orestias* | Opostegidae | 0.52 |  | 53792870 |
| *Opostega sp. CRS1* | Opostegidae | 0.52 |  | 55900069 |
| *Opostega sp. CRS2* | Opostegidae | 0.52 |  | 56618537 |
| *Opostega sp. CRS3* | Opostegidae | 0.52 |  | 52434121 |
| *Opostega sp. CRS4* | Opostegidae | incidental |  | 58100714 |
| *Opostega sp. CRS5* | Opostegidae | 0.52 |  | 53555326 |
| *Opostega sp. CRS6* | Opostegidae | 0.52 |  | 51554193 |
| *Opostega sp. CRS7* | Opostegidae | 0.52 |  | 56583719 |
| *Opostegidae gensp. CRS1* | Opostegidae | 0.52 |  | 51554192 |
| *Azaleodes fuscipes* | Palaephatidae | 1.05 |  | 53207428 |
| *Leuroperna sera* | Plutellidae | 0.52 |  | 49307834 |
| *Plutella xylostella-australiana* | Plutellidae | 8.38 |  | 56928143 |
| *Conoeca guildingi* | Psychidae | 0.52 | Yes | 56923826 |
| *Lepidoscia sp. 10 in BOLD* | Psychidae | 6.81 |  | 31844752 |
| *Psychidae gensp. CRS1* | Psychidae | 0.52 |  | 51349607 |
| *Psychidae gensp. CRS2* | Psychidae | 0.52 |  | 49808173 |
| *Psychidae gensp. CRS3* | Psychidae | 1.05 |  | 51856676 |
| *Lantanophaga pusillidactylus* | Pterophoridae | 1.05 |  | 55861694 |
| *Megalorhipida leucodactylus* | Pterophoridae | 2.62 |  | 30918307 |
| *Pterophoridae gensp. CRS1* | Pterophoridae | 0.52 |  | 50023766 |
| *Pterophoridae gensp. CRS2* | Pterophoridae | 0.52 |  | 31844792 |
| *Pterophoridae gensp. CRS3* | Pterophoridae | 4.19 |  | 49097034 |
| *Pterophoridae gensp. CRS4* | Pterophoridae | 0.52 |  | 55377944 |
| *Sphenarches anisodactylus* | Pterophoridae | incidental |  | 60185508 |
| *Stangeia xerodes* | Pterophoridae | 1.57 |  | 30806623 |
| *Stenoptilodes taprobanes* | Pterophoridae | 3.14 |  | 31605654 |
| *Arescoptera idiotypa* | Pyralidae | 0.52 |  | 55980649 |
| *Assara seminivale* | Pyralidae | 0.52 |  | 49907493 |
| *Assara subarcuella* | Pyralidae | 2.62 |  | 52440197 |
| *Aurana actiosella* | Pyralidae | 1.57 |  | 54707531 |
| *Conobathra hemichlaena* | Pyralidae | 1.05 |  | 56289561 |
| *Curena caustopa* | Pyralidae | 8.9 |  | 47382252 |
| *Doloessa viridis* | Pyralidae | 1.57 |  | 49808267 |
| *Endotricha approximalis* | Pyralidae | 7.85 |  | 49829855 |
| *Endotricha ignealis* | Pyralidae | 1.05 |  | 55377954 |
| *Endotricha lobibasalis* | Pyralidae | 1.57 |  | 56618517 |
| *Endotricha mesenterialis* | Pyralidae | 23.04 |  | 52924434 |
| *Ephestiopsis oenobarella* | Pyralidae | 0.52 |  | 56327471 |
| *Epicrocis metallopa* | Pyralidae | 0.52 |  | 54886172 |
| *Epicrocis pulchra* | Pyralidae | incidental |  | 57098105 |
| *Epipaschiinae gensp. CRS1* | Pyralidae | 1.57 |  | 52204497 |
| *Epipaschiinae gensp. CRS2* | Pyralidae | 0.52 |  | 56327484 |
| *Epipaschiinae gensp. CRS3* | Pyralidae | 0.52 |  | 52345700 |
| *Etiella chrysoporella* | Pyralidae | 0.52 |  | 56319197 |
| *Etiella sp. CRS1* | Pyralidae | 1.57 |  | 50671003 |
| *Etiella sp. CRS2* | Pyralidae | 1.05 |  | 49085738 |
| *Etiella walsinghamella* | Pyralidae | 1.05 |  | 50023789 |
| *Etiella zinckenella* | Pyralidae | 0.52 |  | 56583743 |
| *Gauna flavibasalis* | Pyralidae | 2.62 |  | 53792889 |
| *Guastica auropurpurella* | Pyralidae | incidental |  | 57102502 |
| *Heteromicta poeodes* | Pyralidae | 0.52 |  | 51172694 |
| *Hypargyria metalliferella* | Pyralidae | 1.05 |  | 55873535 |
| *Lacalma albirufalis* | Pyralidae | 0.52 |  | 45858859 |
| *Lamoria eumeces* | Pyralidae | 0.52 |  | 51856669 |
| *Mampava rhodoneura* | Pyralidae | 0.52 |  | 50036610 |
| *Medaniaria adiacritis* | Pyralidae | 2.09 |  | 55692147 |
| *Morosaphycita oculiferella* | Pyralidae | 1.57 |  | 54694850 |
| *Morosaphycita tridens* | Pyralidae | 0.52 |  | 56023520 |
| *Orthaga sp. CRS1* | Pyralidae | 0.52 |  | 54707542 |
| *Orthaga sp. CRS2* | Pyralidae | 0.52 |  | 53792896 |
| *Phycitinae aff. Epicrocis* | Pyralidae | 0.52 |  | 50124318 |
| *Phycitinae aff. Etiella* | Pyralidae | incidental |  | 58563319 |
| *Phycitinae gensp. CRS1* | Pyralidae | 1.05 |  | 53247983 |
| *Phycitinae gensp. CRS2* | Pyralidae | 0.52 |  | 53246576 |
| *Phycitinae gensp. CRS3* | Pyralidae | 3.66 |  | 52927259 |
| *Phycitinae gensp. CRS4* | Pyralidae | 0.52 |  | 52724303 |
| *Phycitinae gensp. CRS5* | Pyralidae | 0.52 |  | 55377953 |
| *Phycitinae gensp. CRS6* | Pyralidae | 0.52 |  | 49808231 |
| *Phycitinae gensp. CRS7* | Pyralidae | 0.52 |  | 49907481 |
| *Phycitinae gensp. CRS8* | Pyralidae | 2.09 |  | 51993215 |
| *Phycitinae gensp. CRS9* | Pyralidae | 3.14 |  | 55466664 |
| *Phycitinae gensp. CRS10* | Pyralidae | 0.52 |  | 56590897 |
| *Phycitinae gensp. CRS11* | Pyralidae | 0.52 |  | 56327485 |
| *Phycitinae gensp. CRS12* | Pyralidae | 1.05 |  | 53035517 |
| *Phycitinae gensp. CRS13* | Pyralidae | 0.52 |  | 54243538 |
| *Phycitinae gensp. CRS14* | Pyralidae | 1.05 |  | 53246595 |
| *Phycitinae gensp. CRS15* | Pyralidae | 0.52 |  | 48898463 |
| *Phycitinae gensp. CRS16* | Pyralidae | 0.52 |  | 54707551 |
| *Phycitinae gensp. CRS17* | Pyralidae | 1.05 |  | 55695051 |
| *Pyralidae gensp. CRS1* | Pyralidae | 0.52 |  | 43305185 |
| *Pyralidae gensp. CRS2* | Pyralidae | 0.52 |  | 49718886 |
| *Pyralidae gensp. CRS3* | Pyralidae | 0.52 |  | 31605677 |
| *Pyralidae GreenMetasia* | Pyralidae | 0.52 |  | 56590908 |
| *Pyralinae Embrace* | Pyralidae | 2.09 |  | 47382194 |
| *Termioptycha eucarta* | Pyralidae | 2.09 |  | 47382280 |
| *Thalamorrhyncha isoneura* | Pyralidae | 1.05 |  | 51669389 |
| *Tirathaba rufivena* | Pyralidae | 0.52 |  | 50773394 |
| *Coscinocera hercules* | Saturniidae | 0.52 |  | 37879469 |
| *Opodiphthera fervida* | Saturniidae | 0.52 |  | 34355149 |
| *Syntherata escarlata* | Saturniidae | 0.52 |  | 43299667 |
| *Syntherata leonae* | Saturniidae | 0.52 |  | 40357121 |
| *Agrius convolvuli* | Sphingidae | 1.05 |  | 38830763 |
| *Cizara ardeniae* | Sphingidae | incidental |  | 41254069 |
| *Gnathothlibus eras* | Sphingidae | 0.52 |  | 49917960 |
| *Hyles livornicoides* | Sphingidae | 1.57 |  | 55980658 |
| *Psilogramma menephron ssp. nebulosa* | Sphingidae | incidental |  | 97523594 |
| *Theretra oldenlandiae* | Sphingidae | 1.05 |  | 39399623 |
| *Theretra turneri* | Sphingidae | incidental |  | 69304614 |
| *Hieromantis ephodophora* | Stathmopodidae | 3.14 |  | 49085795 |
| *Stathmopoda aff. crocophanes* | Stathmopodidae | incidental |  | 58563315 |
| *Stathmopoda castanodes* | Stathmopodidae | 5.24 |  | 51554202 |
| *Stathmopoda megathyma* | Stathmopodidae | 1.05 |  | 56289628 |
| *Stathmopoda melanochra* | Stathmopodidae | 0.52 |  | 47382166 |
| *Stathmopoda triselena* | Stathmopodidae | 0.52 |  | 49808233 |
| *Stathmopoda WhiteThorax* | Stathmopodidae | incidental |  | 57095065 |
| *Stathmopoda sp. DarkOrangeFudge* | Stathmopodidae | 1.05 |  | 54499922 |
| *Stathmopoda sp. Goldwings* | Stathmopodidae | 0.52 |  | 52879184 |
| *Stathmopoda sp. OrangeThreeBrown* | Stathmopodidae | 0.52 |  | 54243547 |
| *Stathmopoda sp. OrangeTwoBrown* | Stathmopodidae | 3.14 |  | 56618439 |
| *Stathmopoda sp. ThreeBrownSpots* | Stathmopodidae | 3.66 |  | 53207462 |
| *Stathmopoda sp. TwoWhiteSpots* | Stathmopodidae | 0.52 |  | 52879176 |
| *Stathmopoda sp. WhiteTwoBrown* | Stathmopodidae | 1.05 |  | 56895229 |
| *Stathmopodidae gensp. CRS1* | Stathmopodidae | 0.52 |  | 49808237 |
| *Stathmopodidae gensp. CRS2* | Stathmopodidae | 0.52 |  | 49085760 |
| *Stathmopodidae gensp. CRS3* | Stathmopodidae | 0.52 |  | 56004714 |
| *Stathmopodidae gensp. CRS4* | Stathmopodidae | 0.52 |  | 56370383 |
| *Stathmopodinae gensp. CRS3* | Stathmopodidae | 0.52 |  | 52724277 |
| *Addaea fragilis* | Thyrididae | 0.52 |  | 54707533 |
| *Canaea hyalospila* | Thyrididae | incidental |  | 68152118 |
| *Hypolamprus bastialis* | Thyrididae | incidental |  | 100135952 |
| *Hypolamprus melilialis* | Thyrididae | 0.52 |  | 50012331 |
| *Mellea ordinaria* | Thyrididae | 0.52 |  | 53792916 |
| *Oxycophina theorina* | Thyrididae | 0.52 |  | 40753547 |
| *Pharambara splendida* | Thyrididae | 1.05 |  | 50202175 |
| *Edosa aff sp. 12 in BOLD* | Tineidae | 1.05 |  | 56299177 |
| *Edosa sp. 7 in BOLD* | Tineidae | 2.09 |  | 49085804 |
| *Edosa xystidophora* | Tineidae | incidental |  | 42199832 |
| *Erechthias sp. CRS1* | Tineidae | incidental |  | 57102505 |
| *Monopis chrysogramma* | Tineidae | 2.62 |  | 31780463 |
| *Monopis crocicapitella* | Tineidae | 1.05 |  | 55362885 |
| *Monopis ethelella* | Tineidae | 1.05 |  | 55989598 |
| *Monopis icterogastra* | Tineidae | 2.62 |  | 31844775 |
| *Niditinea fuscella* | Tineidae | 0.52 |  | 55900063 |
| *Opogona stenocraspeda* | Tineidae | 3.66 |  | 54707582 |
| *Praeacedes atomosella* | Tineidae | 10.99 |  | 56618488 |
| *Tineidae gensp. CRS1* | Tineidae | incidental |  | 59928451 |
| *Tineiodea gensp. CRS1* | Tineidae | incidental |  | 59314697 |
| *Euthrausta phoenicea* | Tineodidae | 1.05 | Yes | 31780471 |
| *Acropolitis canana* | Tortricidae | 0.52 |  | 56928149 |
| *Adoxophyes aff. moderatana* | Tortricidae | 0.52 |  | 54707476 |
| *Adoxophyes sp. C in ANIC* | Tortricidae | 0.52 |  | 49307829 |
| *Adoxophyes sp. D in ANIC* | Tortricidae | 1.57 |  | 53324915 |
| *Anisogona simana* | Tortricidae | 0.52 |  | 55135120 |
| *Coeloptera gyrobathra* | Tortricidae | 0.52 |  | 56466035 |
| *Crocidosema lantana* | Tortricidae | 0.52 |  | 31152529 |
| *Crocidosema plebejana* | Tortricidae | 5.76 |  | 54907847 |
| *Cryptaspasma sordida* | Tortricidae | 2.09 |  | 55362865 |
| *Cryptoptila australana* | Tortricidae | 8.9 |  | 49723542 |
| *Epiphyas aff. sobrina* | Tortricidae | 0.52 |  | 56683785 |
| *Epiphyas sp. CRS1* | Tortricidae | 0.52 |  | 53016855 |
| *Epitymbia eudrosa* | Tortricidae | 1.05 |  | 56583733 |
| *Holocola aff. deloschema* | Tortricidae | 0.52 |  | 56583740 |
| *Isotenes miserana* | Tortricidae | 0.52 |  | 49808247 |
| *Olethreutinae aff. Crocidosema* | Tortricidae | 0.52 |  | 56327488 |
| *Olethreutinae Divergenttips* | Tortricidae | 1.57 |  | 56928167 |
| *Olethreutinae gensp. CRS1* | Tortricidae | 0.52 |  | 55900035 |
| *Olethreutinae gensp. CRS2* | Tortricidae | 0.52 |  | 56618473 |
| *Olethreutinae gensp. CRS3* | Tortricidae | 0.52 |  | 56618501 |
| *Olethreutinae gensp. CRS4* | Tortricidae | 0.52 |  | 56916532 |
| *Olethreutinae gensp. CRS5* | Tortricidae | incidental |  | 58480360 |
| *Olethreutinae GreyBrownPetticoat* | Tortricidae | 0.52 |  | 56618466 |
| *Olethreutinae Puglike* | Tortricidae | 0.52 |  | 54522296 |
| *Phricanthes asperana* | Tortricidae | 0.52 |  | 56928141 |
| *Podognatha vinculata* | Tortricidae | 0.52 |  | 49202158 |
| *Thaumatotibia aclyta* | Tortricidae | 0.52 |  | 54907882 |
| *Tortricidae BlackTuft* | Tortricidae | 0.52 |  | 49808270 |
| *Tortricidae gensp. CRS1* | Tortricidae | 0.52 |  | 31464709 |
| *Tortricidae gensp. CRS2* | Tortricidae | 0.52 |  | 52999336 |
| *Tortricidae gensp. CRS3* | Tortricidae | 0.52 |  | 52879172 |
| *Tortricidae gensp. CRS4* | Tortricidae | 0.52 |  | 55989591 |
| *Tortricidae gensp. CRS5* | Tortricidae | 0.52 |  | 52434114 |
| *Tortricidae gensp. CRS6* | Tortricidae | 0.52 |  | 54495190 |
| *Tortricidae IncaMarkings* | Tortricidae | 1.57 |  | 31780450 |
| *Tortricidae KneelingMan* | Tortricidae | 1.05 |  | 51569172 |
| *Tortricidae PureGold* | Tortricidae | 0.52 |  | 31096108 |
| *Tortricidae Shoulderpads* | Tortricidae | 0.52 |  | 54907884 |
| *Tortricidae SilverScales* | Tortricidae | 0.52 |  | 56466054 |
| *Tortricidae SilverSpangled* | Tortricidae | 1.57 |  | 55692149 |
| *Tortricidae SkeletonKing* | Tortricidae | 0.52 |  | 52554677 |
| *Tortricidae SomeSilverSplangles* | Tortricidae | 1.05 |  | 52435467 |
| *Tortricinae aff Capnoptycha zostrophora* | Tortricidae | incidental |  | 58100712 |
| *Tortricinae aff. Epiphyas* | Tortricidae | 0.52 |  | 51854393 |
| *Tortricinae gensp. CRS1* | Tortricidae | incidental |  | 58563320 |
| *Tortricinae gensp. CRS2* | Tortricidae | incidental |  | 59928446 |
| *Trymalitis sp. ANIC 1 in BOLD* | Tortricidae | 0.52 |  | 47382275 |
| *Alcides metaurus* | Uraniidae | 0.52 |  | 47464472 |
| *Chundana lugubris* | Uraniidae | 2.09 |  | 50343382 |
| *Dirades lugens* | Uraniidae | 0.52 |  | 55900070 |
| *Epiplema coeruleotincta* | Uraniidae | 2.09 |  | 50106817 |
| *Epiplema desistaria* | Uraniidae | 0.52 |  | 49085777 |
| *Epiplema quadristrigata* | Uraniidae | 0.52 |  | 50639101 |
| *Epipleminae gensp. CRS1* | Uraniidae | 0.52 |  | 50338652 |
| *Micronia aculeata* | Uraniidae | incidental |  | 97523601 |
| *Phazaca leucocephala* | Uraniidae | 1.05 |  | 49723585 |
| *Phazaca mutans* | Uraniidae | 3.14 |  | 47382200 |
| *Phazaca sp. CRS1* | Uraniidae | 0.52 |  | 49307848 |
| *Phazaca sp. CRS2* | Uraniidae | 1.05 |  | 54707528 |
| *Phazaca stolida* | Uraniidae | 0.52 |  | 48898399 |
| *Arignota stercorata* | Xyloryctidae | incidental |  | 60059605 |
| *Catoryctis eugramma* | Xyloryctidae | 0.52 |  | 56370382 |
| *Echiomima mythica* | Xyloryctidae | incidental |  | 62842278 |
| *Iulactis sp. CRS1* | Xyloryctidae | incidental |  | 61786599 |
| *Maroga melanostigma* | Xyloryctidae | 1.57 |  | 49718883 |
| *Xylorycta luteotactella* | Xyloryctidae | 0.52 |  | 33125018 |
| *Chionogenes trimetra* | Yponomeutidae | incidental |  | 61473070 |
| *Niphonympha oxydelta* | Yponomeutidae | 1.05 |  | 56004668 |
| *Yponomeuta paurodes* | Yponomeutidae | 0.52 |  | 31096105 |
| *Yponomeutidae gensp. CRS1* | Yponomeutidae | 0.52 |  | 56327482 |
| *Yponomeutidae gensp. CRS2* | Yponomeutidae | 1.05 |  | 55362904 |
| *Yponomeutidae gensp. CRS3* | Yponomeutidae | 13.61 |  | 56295621 |
| *Pollanisus sp. CRS1* | Zygaenidae | 1.57 |  | 31844764 |
| *Gelechioidea aff Barea* |  | incidental |  | 58480284 |
| *Gelechioidea aff Blastobasis* |  | incidental |  | 58574171 |
| *Gelechioidea aff Chezala* |  | 0.52 |  | 54694862 |
| *Gelechioidea aff Eulechria* |  | 0.52 |  | 54707594 |
| *Gelechioidea aff Thiotricha* |  | 0.52 |  | 56728660 |
| *Gelechioidea Blackhead* |  | 6.81 |  | 55900038 |
| *Gelechioidea BrownYellowBrown* |  | incidental |  | 58563321 |
| *Gelechioidea gensp. CRS1* |  | 0.52 |  | 54367874 |
| *Gelechioidea gensp. CRS2* |  | 0.52 |  | 54495202 |
| *Gelechioidea gensp. CRS3* |  | 0.52 |  | 56583638 |
| *Gelechioidea gensp. CRS4* |  | 0.52 |  | 58010821 |
| *Gelechioidea gensp. CRS5* |  | 0.52 |  | 56618513 |
| *Gelechioidea gensp. CRS6* |  | 0.52 |  | 54694871 |
| *Gelechioidea gensp. CRS7* |  | 0.52 |  | 54707525 |
| *Gelechioidea gensp. CRS8* |  | 0.52 |  | 56370365 |
| *Gelechioidea gensp. CRS9* |  | 0.52 |  | 49085703 |
| *Gelechioidea gensp. CRS10* |  | 0.52 |  | 33125061 |
| *Gelechioidea gensp. CRS11* |  | 0.52 |  | 56618504 |
| *Gelechioidea gensp. CRS12* |  | 4.19 |  | 52879218 |
| *Gelechioidea gensp. CRS 13* |  | 1.05 |  | 55224286 |
| *Gelechioidea gensp. CRS14* |  | 0.52 |  | 52879215 |
| *Gelechioidea gensp. CRS15* |  | 0.52 |  | 52440207 |
| *Gelechioidea gensp. CRS16* |  | 0.52 |  | 55224196 |
| *Gelechioidea gensp. CRS 17* |  | 0.52 |  | 31096121 |
| *Gelechioidea gensp. CRS18* |  | incidental |  | 60024757 |
| *Gelechioidea gensp. CRS19* |  | incidental |  | 60190445 |
| *Gelechioidea gensp. CRS20* |  | incidental |  | 60312310 |
| *Gelechioidea gensp. CRS21* |  | incidental |  | 59419158 |
| *Gelechioidea gensp. Golden* |  | 1.57 |  | 54907798 |
| *Gelechioidea gensp. RustyOrange* |  | 0.52 |  | 56728608 |
| *Gelechioidea Grey Philobota* |  | incidental |  | 58480358 |
| *Gelechioidea Spotty* |  | 0.52 |  | 56979922 |
| *Gelechioidea Zebra* |  | 0.52 |  | 33125042 |
| *Gracillarioidea aff. Phyllocnistis* |  | 0.52 |  | 55224194 |
| *Gracillarioidea gensp. 1* |  | 0.52 |  | 54694841 |
| *Lepidoptera aff Xylorctidae CRS1* |  | 1.05 |  | 50004074 |
| *Lepidoptera aff. Cosmopterigidae* |  | 0.52 |  | 56466063 |
| *Lepidoptera aff. Lecithoceridae* |  | 0.52 |  | 56466055 |
| *Lepidoptera aff. Opogona* |  | incidental |  | 59108464 |
| *Lepidoptera aff. Opostega* |  | 0.52 |  | 56583738 |
| *Lepidoptera aff. Phytomeria* |  | 0.52 |  | 56618507 |
| *Lepidoptera aff. Scoparinae* |  | 1.57 |  | 33125071 |
| *Lepidoptera gensp. CRS1* |  | 0.52 |  | 55873545 |
| *Lepidoptera gensp. CRS2* |  | 0.52 |  | 49907492 |
| *Lepidoptera gensp. CRS3* |  | 0.52 |  | 54228787 |
| *Lepidoptera gensp. CRS4* |  | 0.52 |  | 48898427 |
| *Lepidoptera gensp. CRS5* |  | 0.52 |  | 56618477 |
| *Lepidoptera gensp. CRS6* |  | 0.52 |  | 56327473 |
| *Lepidoptera gensp. CRS7* |  | 0.52 |  | 56916522 |
| *Lepidoptera gensp. CRS8* |  | 0.52 |  | 56683796 |
| *Lepidoptera gensp. CRS9* |  | 0.52 |  | 56683791 |
| *Lepidoptera gensp. CRS10* |  | 0.52 |  | 56618506 |
| *Lepidoptera gensp. CRS11* |  | 0.52 |  | 56370387 |
| *Lepidoptera gensp. CRS12* |  | 0.52 |  | 56370378 |
| *Lepidoptera gensp. CRS13* |  | 0.52 |  | 56370376 |
| *Lepidoptera gensp. CRS14* |  | 0.52 |  | 56370369 |
| *Lepidoptera gensp. CRS15* |  | 0.52 |  | 56299164 |
| *Lepidoptera gensp. CRS16* |  | 0.52 |  | 51554188 |
| *Lepidoptera gensp. CRS17* |  | 0.52 |  | 53792893 |
| *Lepidoptera gensp. CRS18* |  | 0.52 |  | 53035514 |
| *Lepidoptera gensp. CRS19* |  | 0.52 |  | 30918328 |
| *Lepidoptera gensp. CRS20* |  | 0.52 |  | 56895212 |
| *Lepidoptera gensp. CRS21* |  | incidental |  | 57095066 |
| *Lepidoptera gensp. CRS22* |  | 0.52 |  | 54022470 |
| *Lepidoptera gensp. CRS23* |  | 0.52 |  | 55362873 |
| *Lepidoptera gensp. CRS24* |  | 0.52 |  | 51993211 |
| *Lepidoptera gensp. CRS25* |  | 0.52 |  | 53555327 |
| *Lepidoptera gensp. CRS26* |  | 0.52 |  | 54694839 |
| *Lepidoptera gensp. CRS27* |  | 0.52 |  | 54907837 |
| *Lepidoptera gensp. CRS28* |  | incidental |  | 58574172 |
| *Lepidoptera gensp. CRS29* |  | incidental |  | 57098491 |
| *Lepidoptera gensp. CRS30* |  | incidental |  | 57102501 |
| *Lepidoptera gensp. CRS31* |  | incidental |  | 58318048 |
| *Lepidoptera gensp. CRS32* |  | incidental |  | 60024756 |
| *Lepidoptera gensp. CRS33* |  | incidental |  | 59419156 |
| *Lepidoptera gensp. CRS34* |  | incidental |  | 60190443 |
| *Lepidoptera gensp. CRS35* |  | incidental |  | 60312299 |
| *Lepidoptera gensp. Magneto* |  | 0.52 |  | 56370368 |
| *Lepidoptera YellowWingWindows* |  | 0.52 |  | 56466073 |
| *Noctuiodea aff. Calathusa* |  | 0.52 |  | 55989622 |
| *Noctuoidea gensp. BrownBlueBulldog* |  | 0.52 |  | 56289627 |
| *Noctuoidea gensp. CRS1* |  | 0.52 |  | 56583693 |
| *Noctuoidea gensp. CRS2* |  | 0.52 |  | 49532765 |
| *Noctuoidea gensp. CRS3* |  | 2.09 |  | 33123747 |
